# Activation of the IKK2-NFκB pathway in VSMCs inhibits calcified vascular stiffness in CKD by reducing the secretion of calcifying extracellular vesicles

**DOI:** 10.1101/2023.07.11.548621

**Authors:** Shinobu Miyazaki-Anzai, Masashi Masuda, Audrey L. Keenan, Yuji Shiozaki, Makoto Miyazaki

## Abstract

IKK2-NFκB pathway mediated-inflammation in vascular smooth muscle cells (VSMCs) has been proposed to be an etiologic factor in medial calcification and stiffness. However, the role of the IKK2-NFκB pathway in medial calcification remains to be elucidated. In this study, we found that CKD induces inflammatory pathways through the local activation of the IKK2-NFκB pathway in VMSCs associated with calcified vascular stiffness. Despite reducing the expression of inflammatory mediators, complete inhibition of the IKK2-NFκB pathway *in vitro* and *in vivo* unexpectedly exacerbated vascular mineralization and stiffness. In contrast, activation of NFκB by SMC-specific IκB deficiency attenuated calcified vascular stiffness in CKD. Inhibition of the IKK2-NFκB pathway induced apoptosis of VSMCs by reducing anti-apoptotic gene expression, whereas activation of NFκB reduced CKD-dependent vascular cell death. In addition, increased calcifying extracellular vesicles through the inhibition of the IKK2-NFκB pathway induced mineralization of VSMCs, which was significantly reduced by blocking cell death. This study reveals that activation of the IKK2-NFκB pathway in VSMCs plays a protective role in CKD-dependent calcified vascular stiffness by reducing the release of apoptotic calcifying extracellular vesicles.

## Introduction

Over half of all deaths among chronic kidney disease (CKD) patients are due to cardiovascular disease (CVD). The risk of CVD mortality in CKD patients is 20-30 times higher than that of the general population(1–4). Growing evidence suggests that this increased risk of CVD mortality is explained by the predisposition of CKD patients to vascular calcification(5–8). Vascular calcification occurs at two distinct sites within the vessel wall: the intima and the media. Intimal calcification, also called atherosclerotic calcification, occurs in the context of atherosclerosis and involves lipids, macrophages, and vascular smooth muscle cells (VSMCs)(9, 10). Medial calcification can exist independently of atherosclerosis and is associated with elastin and VSMCs, and is more prevalent in patients with CKD. In both cases, accumulation of calcium-phosphate complexes in the vascular wall decreases aortic elasticity and flexibility, which impairs cardiovascular hemodynamics, resulting in substantial morbidity and mortality(7, 11). CKD is represented by states of low-grade chronic inflammation characterized by increased levels of inflammatory markers such as tumor necrosis factor-a (TNFα) and interleukins (IL)(7, 12). A number of previous studies show that inflammatory cytokines play causative roles in vascular calcification. Treatment with anti-inflammatory agents such anti-TNFα and anti-interleukin (IL-1β and IL-6) monoclonal antibodies have been shown to reduce vascular calcification(13–16). However, the molecular mechanisms by which CKD increases levels of inflammatory factors such as TNFα and ILs locally in VSMCs, resulting in vascular calcification, are not fully investigated.

NFκΒ proteins comprise a family of structurally related eukaryotic transcription factors that are involved in the control of a large number of normal cellular and organismal processes, such as immune and inflammatory responses(17–19). These transcription factors are highly active in a number of disease states including cancer, metabolic diseases (e.g., diabetes and obesity), asthma, neurodegenerative diseases, and cardiovascular diseases(18, 20–24). In most cells, NFκΒ is present as a latent, inactive, IκB-bound complex in the cytosol. When a cell receives any of a multitude of extracellular signals such as LPS and TNFα, NFκΒ rapidly enters the nucleus and activates gene expression. Therefore, a crucial step for regulating NFκΒ activity is the regulation of the IκΒ/NFκΒ interaction. Signals that activate NFκΒ converge on the activation of a regulatory complex that contains a serine-specific IκB kinase (IKK). IKK is an unusual kinase and contains three distinct subunits: IKK1, IKK2 and IKK3. IKK1 and IKK2 compose of catalytic kinase subunits while IKK3 is a regulatory subunit that serves as a sensing scaffold and integrator of upstream signals for activation of the catalytic subunits. In the canonical pathway, activation of the IKK complex leads to phosphorylation by IKK2 of two specific serines near the N terminus of IκΒα, which targets IκΒα for ubiquitination and degradation by the 26S proteasome. The released NFκΒ complex can then enter the nucleus to activate the expression of target inflammatory cytokines and chemokine genes(18, 19). Several in vitro and in vivo studies have proposed that NFκB activation contributes to the etiology of vascular calcification(25–30). However, the VSMC-specific role in the IKK2-NFκB inflammatory cascade in the regulation of CKD-induced medial calcification has not been fully elucidated.

In addition to inflammation, the IKK2-NFκB pathway governs cell survival by inhibiting apoptosis, which is a major form of the programmed cell death process involving multiple caspase reactions(31–34). NFκB exerts pro-survival effects by inducing the transcription of several anti-apoptotic genes such as cIAPs, XIAP, cFLIP and Bcl2 family members(32, 35, 36). In addition, IKK2 directly inhibits apoptosis by phosphorylating the major pro-apoptotic factor BAD. Previous studies have shown that apoptosis is an early important event in vascular calcification(37–42). Apoptotic VSMCs disassemble and generate subcellular membrane-bound extracellular vesicles called apoptotic bodies (ApoBD, generally 1-5 um in diameter)(40, 43). The formation of ApoBD has been proposed to play a critical role in vascular mineralization. In addition, recent studies revealed that apoptosis induces the release of smaller extracellular microvesicles (EV) such as apoptotic microvesicles (0.5-0.2 μm) and exosomes (0.2-0.1 μm)(44–46). In addition, recent evidence on the mechanisms of vascular calcification identified calcifying EV-containing mineralization inducers such as alkaline phosphatase (ALP) derived from VSMCs as the mediators of cardiovascular mineralization(47–50). In this study, we investigated whether CKD induces IKK2-NFκB–mediated inflammation locally in VSMCs. We examined how the IKK2-NFκB pathway is involved in the mineralization of VMSCs in vitro using the CRISPR/Cas9 system. We also explored whether alterations of the IKK2-NFκB pathway affect vascular calcification in CKD mouse models.

## Results

CKD is represented by states of low-grade chronic inflammation characterized by increased systemic levels of inflammatory markers such as TNFα and interleukins(51, 52). In addition, numerous studies have indicated that local IKK2-NFκΒ-mediated inflammation in VSMCs plays a causative role in regulating vascular calcification(25–30). Consistent with our previous studies(14, 38, 39, 42, 53, 54), 5/6 nephrectomized DBA SMMHC-GFP mice had significantly higher levels of serum creatinine compared to sham-operated mice (mice with normal kidney function, NKD), which indicates that 5/6 nephrectomy induces CKD. CKD but not NKD SMMHC-GFP mice developed medial calcification (Supplemental Figure 1A-1C). In addition, levels of aortic calcium were >5-fold higher in CKD mice (Supplemental Figure 1D). To determine whether CKD induced vascular stiffness along with vascular calcification, aortic pulse wave velocity was analyzed with an Indus Doppler Flow Velocity System. The aortic pulse wave velocity was greater in CKD compared with NKD mice (Supplemental Figure 1E and 1F). These results indicate that CKD induces calcified artery stiffening. To examine whether CKD induces inflammatory signals locally in VSMCs, we performed RNA-seq using cell-sorting on VSMCs from the aortas of CKD and NKD mice. As shown in Figure 1A, the mRNA-seq and pathway analyses confirmed that genes in numerous inflammatory pathways were increased in the VSMCs of CKD mice compared to NKD mice. At the top of the list, levels of 116 genes involved in the inflammatory response were significantly increased in the VSMCs isolated from CKD mice (Figure 1B). EMSA analysis showed that the NFκB pathway was drastically activated in the VSMCs of CKD mice. In addition, levels of active phosphorylated IKK2 (p-IKK2) and p65 (p-p65) as well as inflammatory markers (IL1β, IL-6, TNFα and iNOS) were higher in the aortic media of CKD mice (Figure 1C-1E). These data suggest that CKD induces IKK2-NFκB-mediated inflammation in VSMCs.

**Figure 1:**
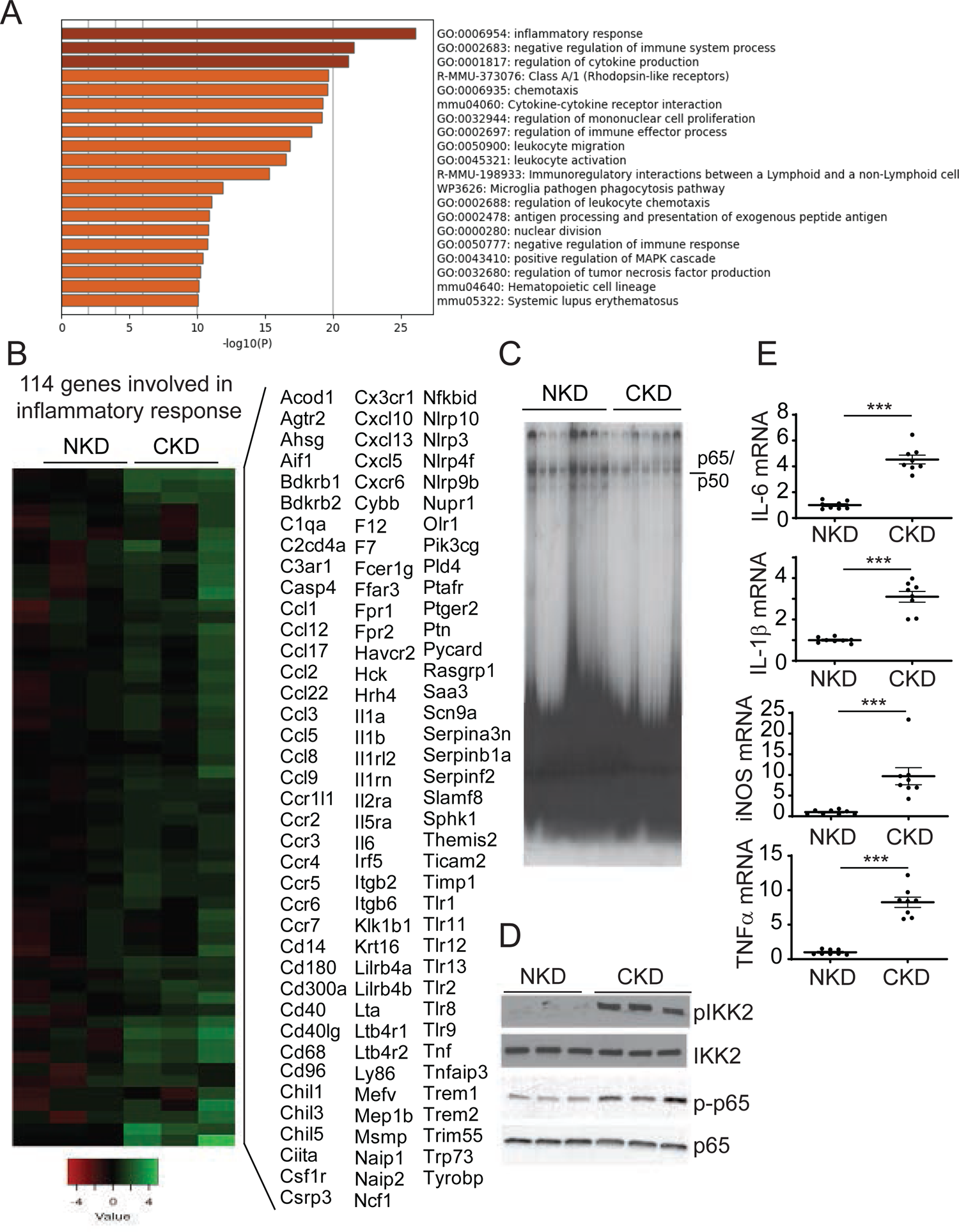
IKK2/NFκB-mediated pro-inflammatory pathways were induced in VSMCs by CKD. A) Pathway analysis of mRNA-seq. Eight-week-old male DBA SMMHC-GFP mice were subjected to either 5/6 nephrectomy (CKD) or sham operation (NKD). Aortas were dissected from CKD and NKD mice 12 weeks after the surgery. SMMHC-GFP+ VSMCs were isolated through cell sorting after the digestion of aortas with collagenase/elastase mixture. Total RNAs were isolated and subjected to mRNA-seq. The mRNA-seq data were deposited as GSE229679. The genes upregulated by CKD were subjected to Metascape pathway analysis (https://metascape.org). B) 116 genes involved in inflammatory response were most strongly induced in the VSMCs of CKD mice. C) Active p65/p50 NFκB complex was increased in the VSMCs from CKD mice. Levels of p65/p50 complex were analyzed by EMSA analysis. D) Immunoblot analysis of the IKK2-NFκB pathway. Total protein lysates were isolated from aortic media. E) Inflammatory marker expression in the aortic media of CKD mice. qPCR analysis confirmed that CKD induced the expression of inflammatory makers in the aortic media. *P<0.05.

CKD induces IKK2-NFκΒ-mediated inflammation locally in VSMCs. Previous in vitro studies conducted to study the role of IKK2-NFκΒ by partial inhibition use chemical inhibitors, dominant negatives, or RNAi (26, 28, 30, 55, 56). To reduce complications from remaining IKK2 activity and off-target effects, we generated IKK2KO VSMCs using a CRISPR-cas9 system (Figure 2A). TNFα treatment induced IKK2-mediated phosphorylation of IκB and p65 in wild-type VSMCs, whereas IKK2 deficiency blocked TNFα-induced phosphorylation of IκB and p65 (Figure 2A). In addition, IKK2 deficiency abolished inflammatory maker inductions by TNFα treatment (Figure 2B) in addition to high-phosphate treatment (Supplemental Figure 2A and 2B). Unexpectedly, however, IKK2 deficiency aggravated high-phosphate and TNFα-induced mineralization (Figure 2C-2E) and osteogenic differentiation of VSMCs (Figure 2F). Similar to CRISPR-cas9-mediated IKK2KO, the overexpression of IKK2 kinase-dead (K33M) dominant negative in VSMCs induced TNFα-induced mineralization of VSMCs (Supplemental Figure 2C and 2D), suggesting that the anti-calcific effect of IKK2 is kinase activity-dependent.

**Figure 2:**
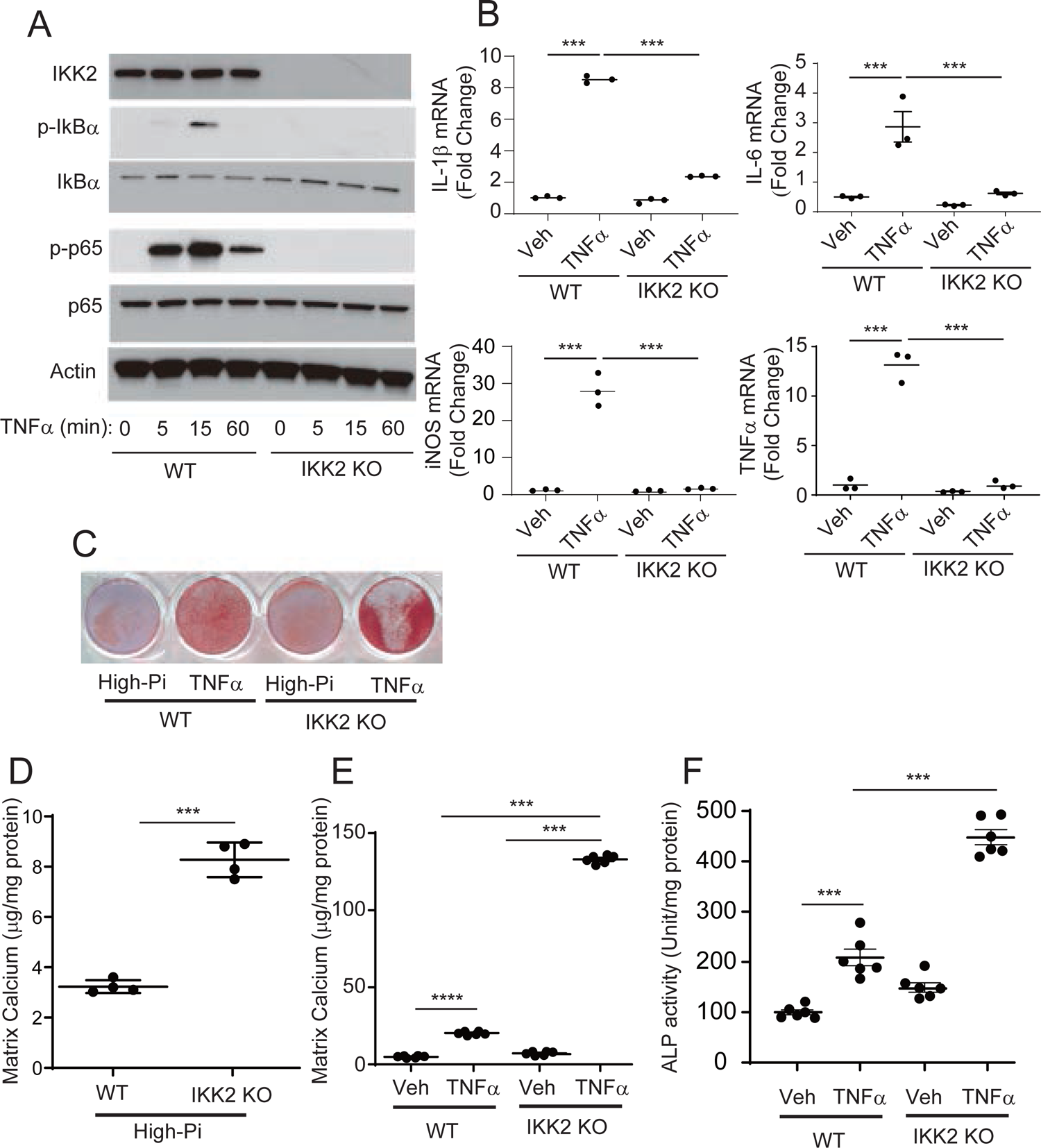
Deletion of the IKK2 gene induces mineralization of VSMCs despite the inhibition of NFκB-mediated inflammatory cytokine induction. A) Immunoblot analysis of the IKK2-NFκB pathway in IKK2KO VSMCs treated with TNFα. B) mRNA levels of inflammatory markers (IL-1β, IL-6, iNOS and TNFα) in IKK2KO VSMCs treated with TNFα. VSMCs were treated with TNFα for 8 hours. Levels of 36B4 mRNA were used as control. C-F) Alizarin Red staining, levels of matrix calcium and ALP activity of VSMCs treated with either high-phosphate or TNFα. VSMCs were treated with either high-phosphate (2.4 mM) or TNFα for 6 days. ***P<0.001

The results from the IKK2 KO VSMCs were unexpected and led us to hypothesize that NFκB-mediated inflammation has distinct effects on vascular calcification from the IKK2 effect. To pursue our hypothesis, we next created VSMCs lacking NFKB1 and RelA genes that produce the p50 and p65 NFκB subunits, respectively (Figure 3A and 3B). Similar to IKK2 deficiency, both loss of p50 and p65 NFκB subunits induced mineralization of VSMCs in response to high-phosphate and TNFα treatment (Figure 3C and 3D). While inducing vascular calcification, p65 deficiency completely reduced levels of inflammatory mediators such as IL-1β, IL-6 and TNFα (Figure 3E-3G).

**Figure 3:**
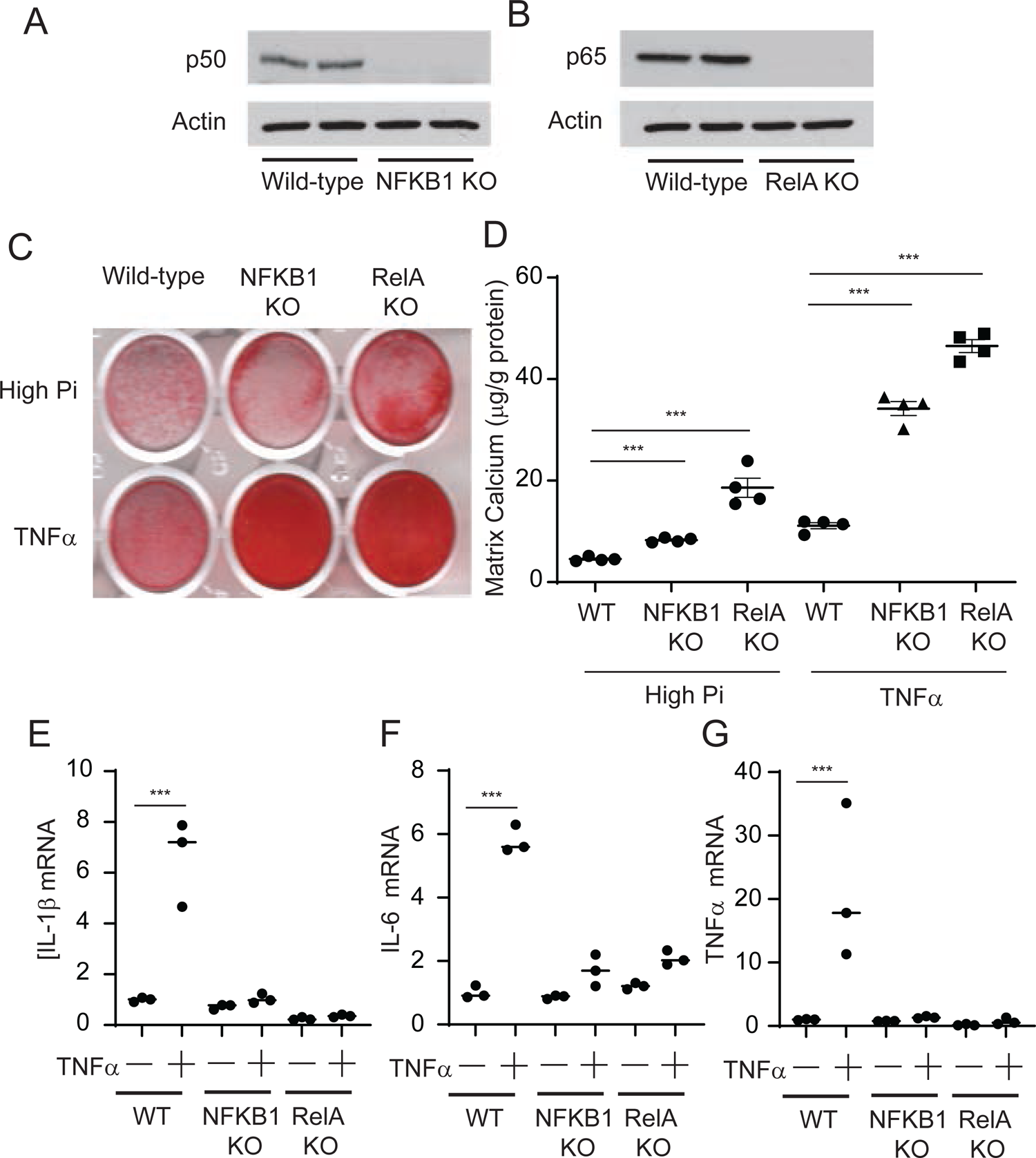
Deletion of NFκB subunits induces mineralization of VSMCs despite the inhibition of NFκB-mediated inflammatory cytokine induction. A, B) Immunoblot analysis of the p50 and p65 NFκB subunits in NFKB1KO and RelAKO VSMCs, respectively. C and D) Alizarin staining, levels of matrix calcium and ALP activity of VSMCs treated with either high-phosphate or TNFα. VSMCs were treated with either high-phosphate (2.4 mM) or TNFα for 6 days. E-G) mRNA levels of inflammatory mediators (IL-1β, IL-6, iNOS and TNFα) in IKK2KO VSMCs treated with TNFα. VSMCs were treated with TNFα for 8 hours. Levels of 36B4 mRNA were used as control. ***P<0.001

To examine whether modulations of the IKK2-NFκB pathway in VSMCs *in vivo* affects CKD-dependent vascular calcification, we generated tamoxifen-inducible VSMC-specific IKK2 knockout (SMC-IKK2KO) mice and subjected them to 5/6 nephrectomy to induce CKD. Tamoxifen injection was able to abolish the expression of IKK2 in the aortic media (Figure 4A) but not adventitia or other tissues (data not shown) of SMC-IKK2KO mice. The significant reduction of IKK2 expression reduced levels of p-IκB and p-p65 in the media that mediate the downstream phosphorylation signaling of IKK2 (Figure 4A). Strikingly, CKD significantly induced early mortality of SMC-IKK2 KO more than control mice (Figure 4B). Because of the CKD-induced early death of SMC-IKK2 KO mice, we examined whether SMC-IKK2 deficiency accelerates CKD-dependent vascular calcification and stiffness at an early time point (3 weeks after CKD induction) when control mice do not develop CKD-dependent cardiovascular complications (Figure 4C-4F). 5/6 nephrectomy increased levels of serum creatinine by 2.3-fold in both control and SMC-IKK2 KO mice whereas levels of serum triglyceride were significantly lower in both control and SMC-IKK2 KO mice with CKD (Supplemental Table 1). Histological analysis of the aortic arches with von Kossa stain revealed that SMC-IKK2 deficiency severely aggravated medial calcification under CKD. Calcified lesions and aortic calcium content were about 150-fold and 20-fold greater in CKD SMC-IKK2 KO mice than CKD control mice (Figure 4D and 4E). In addition, the aortic pulse wave velocity was greater in CKD SMC-IKK2 KO mice compared with other groups (Figure 4F). qPCR analysis showed that CKD-induced inflammatory mediators such as IL-1β, IL-6 and TNFα were normalized by SMC-IKK2 deficiency (Figure 4G-4I). We have previously shown that CKD induces vascular cell death. SMC-IKK2 deficiency remarkably induced cell death in the aortic media (Figure 4J-4K).

**Figure 4.**
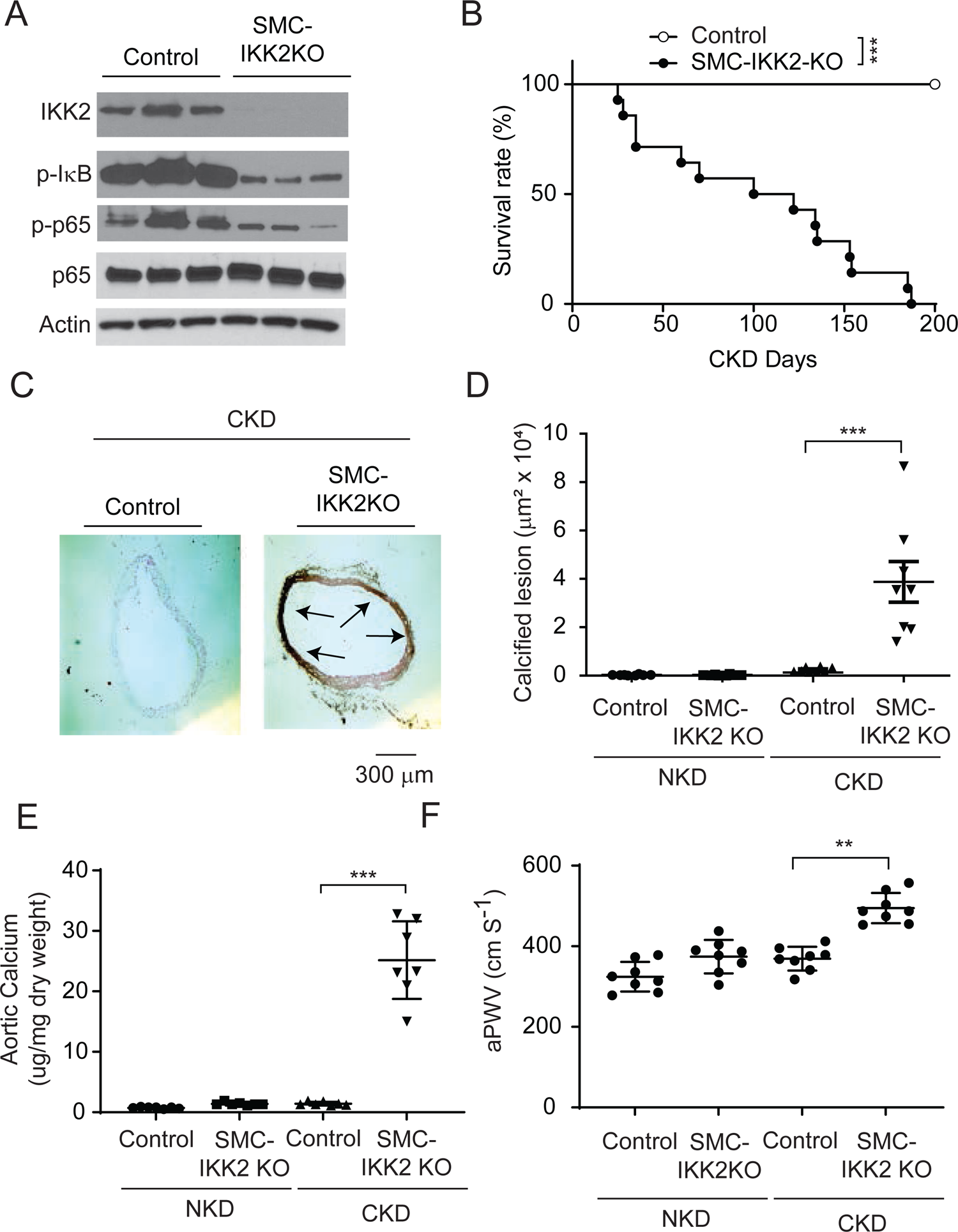
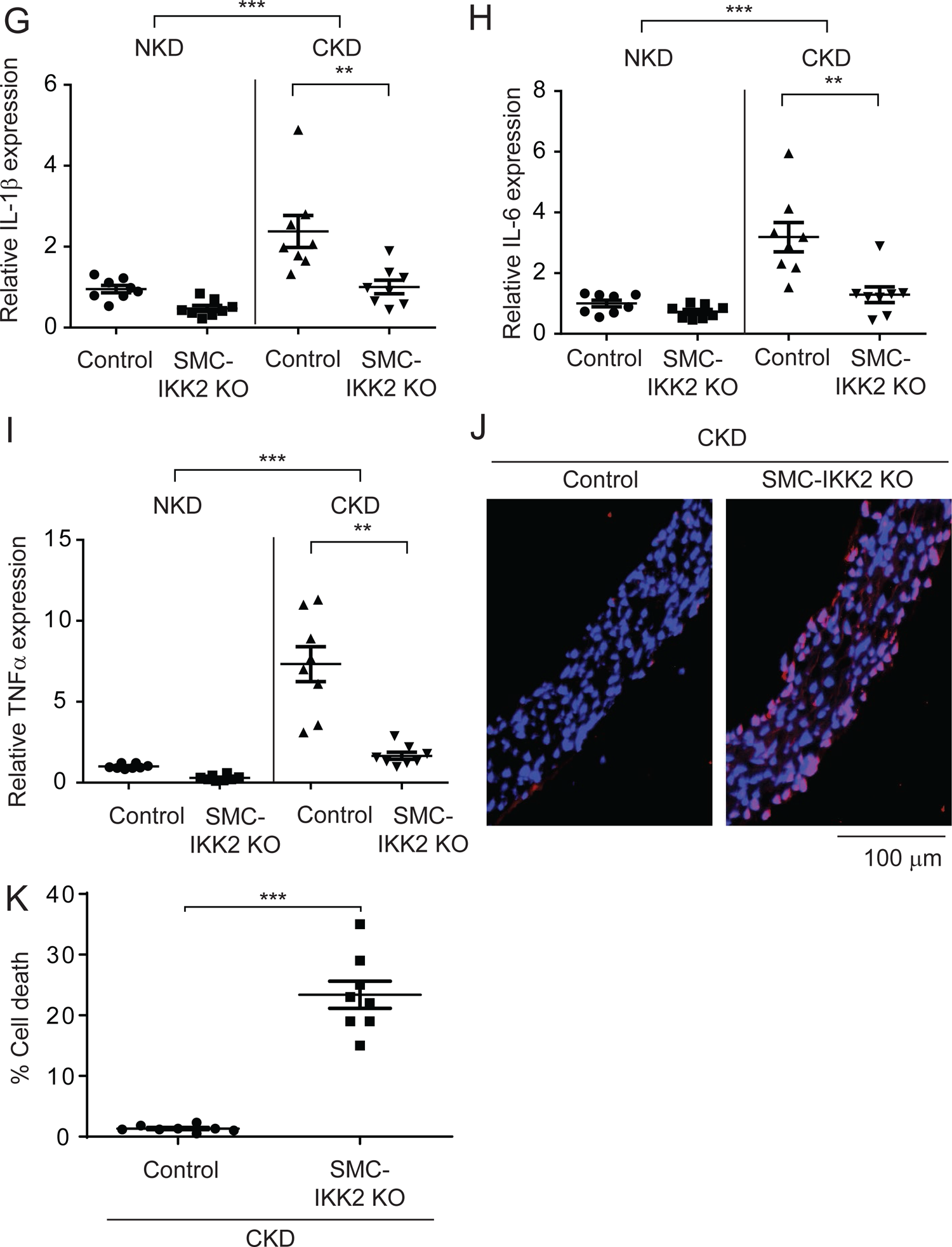
VSMC-IKK2 deficiency induces early mortality and calcified vascular stiffness of mice in CKD. A) Immunoblot analysis of the IKK2-NFκB pathway in the aortic media of SMC-IKK2KO mice. Eight-week-old control and SMC-IKK2KO mice (N=8) were intraperitoneally injected with tamoxifen for 5 days and subjected to either sham-operation (NKD) or 5/6 nephrectomy (CKD). 3 weeks after the surgeries, animals were euthanized. B) Survival rate of SMC-IKK2 mice under CKD. C and D) Histological analysis of aortic arches with von Kossa staining. Aortas were dissected from the mice 3 weeks after CKD was induced. Arrows (black) indicate calcified lesions. E) Aortic calcium content. Aortic calcium content was analyzed with ash assay coupled with calcium colorimetric assay. F) aPWV. aPWV was analyzed using an Indus Doppler Flow Velocity System 3 weeks after CKD was induced. G-I) mRNA levels of inflammatory markers (G, IL-1β, H, IL-6 and I TNFα in the aortic media of SMC-IKK2 mice under NKD and CKD. Aortic media were dissected from the mice 3 weeks after CKD was induced. J and K) Aortic cell death: Cell death was analyzed with a Roche in situ cell death kit. **P<0.01 and ***P<0.001

Since inhibition of the IKK2-NFκB pathway unexpectedly worsened CKD-dependent vascular complications in *in vitro* and *in vivo* models, we next examined how activation of the NFκB pathway affects CKD-dependent vascular complications by knocking out IκBα, which is a central inhibitor of the NFκB pathway. Since global IκBα KO mice were embryonic lethal, we generated SMC-specific IκB KO mice by inserting two loxP sites into intron 1 and intron 5 and crossing with tamoxifen-inducible SMMHC-creER^T2^ (Supplemental Figure 3). Similar to SMC-IKK2 deficiency, tamoxifen injections completely removed IκBα from the aortic media of SMC-IκB KO mice but not control mice, resulting in a significant increase in an active NFκB subunit, p-p65 (Figure 5A). Unlike SMC-IKK2 KO mice, CKD did not affect the mortality of SMC-IκB KO mice (data not shown). The mice were therefore euthanized 12 weeks after CKD was induced, when the control mice develop CKD-dependent vascular complications. CKD increased levels of serum phosphorus and creatinine, while levels of serum triglycerides and calcium were reduced in both control and SMC-IκB KO mice (Supplemental Table 2). Under CKD, SMC-specific IκB deficiency significantly attenuated vascular calcification, stiffness and cell death (Figure 5B-5G). As shown in Figure 1, CKD induced116 inflammatory markers. We next examined whether these CKD-induced inflammatory markers were induced by SMC-IκB deficiency and reduced by SMC-IKK2 deficiency. SMC-IκB deficiency significantly induced about 90% of the CKD-induced inflammatory markers, whereas SMC-IKK2 deficiency reduced about 50% of the CKD-induced inflammatory markers (Figure 5H and Supplemental Table 3).

**Figure 5.**
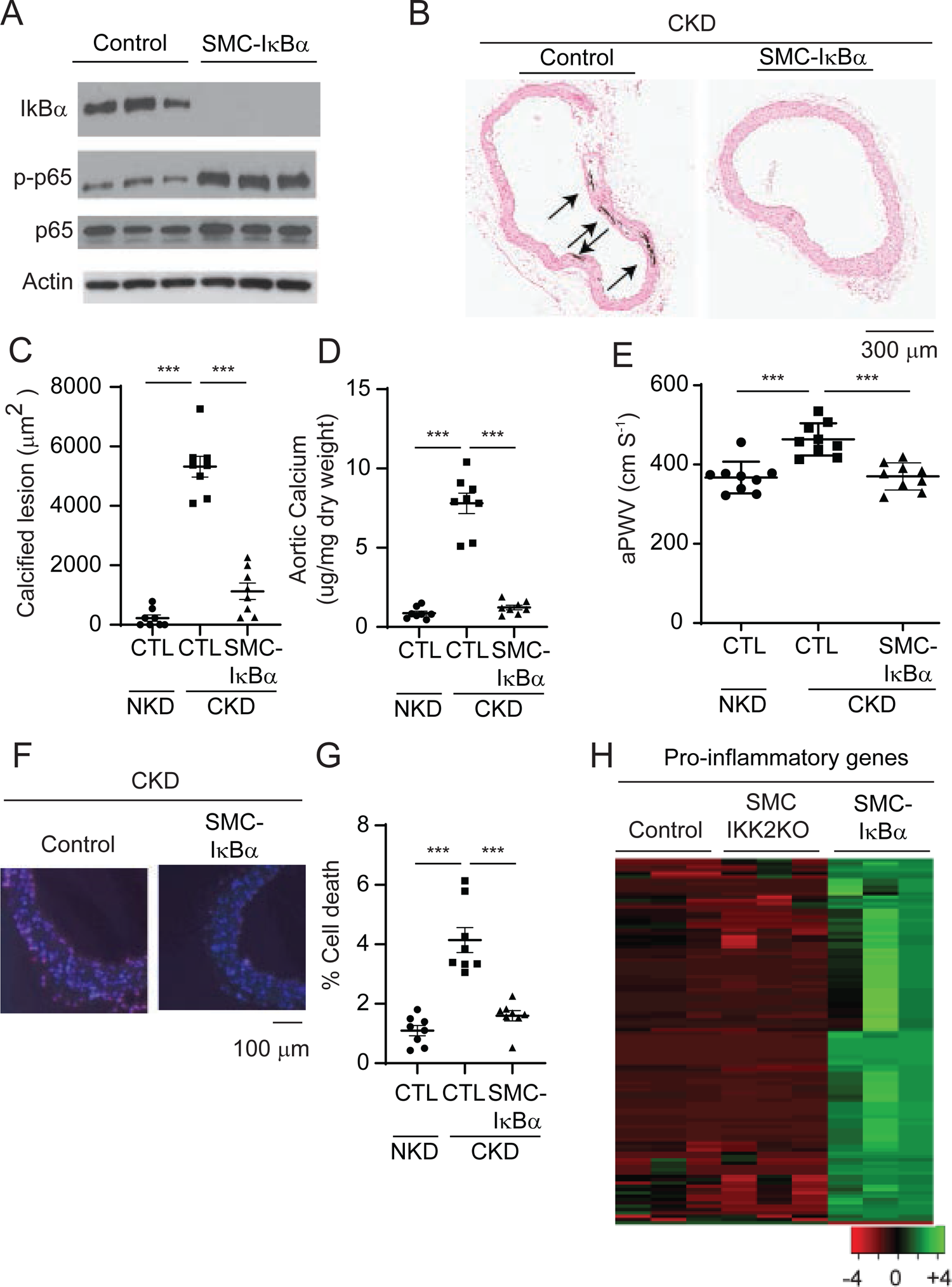
VSMC-IκB deficiency attenuates CKD-induced calcified vascular stiffness. A) Immunoblot analysis of the IKK2-NFκB pathway in the aortic media of SMC-IκBKO mice. Eight-week-old control and SMC-IKK2KO mice (N=8) were intraperitoneally injected with tamoxifen for 5 days and subjected to 5/6 nephrectomy (CKD). 12 weeks after the surgeries, animals were euthanized. B, C) Histological analysis of aortic arches with von Kossa staining. Aortas were dissected from the mice 12 weeks after CKD was induced. Arrows (black) indicate calcified lesions. D) Aortic calcium content. Aortic calcium content was analyzed with ash assay coupled with calcium colorimetric assay. E) aPWV. aPWV was analyzed using an Indus Doppler Flow Velocity System 12 weeks after CKD was induced. F, G) Aortic cell death: Cell death was analyzed with a Roche in situ cell death kit. Aortic media were dissected from the mice 3 weeks after CKD was induced. Pink indicates cell death. H) Heatmap with mRNA levels of >100 inflammatory genes induced by CKD in the aortic media of SMC-IKK2KO and SMC-IκBKO mice. Aortic medial layers were dissected 3 weeks after CKD was induced. The numbers are shown in Supplemental Table 3. ***P<0.001

Severe vascular cell death was associated with CKD-induced and SMC-IKK2 deficiency-induced vascular calcification. The IKK2-NFκB pathway is a critical inhibitory regulator of apoptotic cell death by inducing anti-apoptotic genes in addition to inflammation(31, 32). We next examined whether the inhibition of the IKK2-NFkB pathway induces apoptotic cell death in VSMCs. IKK2 and RelA deficiency enhanced TNFα-induced apoptosis (Figure 6A-6C). TNFα treatment time-dependently activated caspase 3 (Casp3) in IKK2 knockout VSMCs more than wild-type VSMCs (Figure 6A). RelA deficiency also enhanced the activation of Casp3 (Figure 6B). Upon TNFα treatment, both IKK2O and RelAKO VSMCs had significantly higher Casp3 activity (Figure 6C). VSMCs lacking IKK2 and RelA genes had significantly lower levels of several anti-apoptotic genes and proteins such as Bcl2, Bcl2a1a, cIAP2 and XIAP (Figure 6D-6H). Consistently, under CKD, SMC-IκBα deficiency induced 21 out of 22 anti-apoptotic genes expressed in VSMCs, whereas SMC-IKK2 deficiency significantly reduced 18 anti-apoptotic genes (Figure 6I and Supplemental Table 4). Recent studies indicated that apoptosis increases the secretion of extracellular vesicles such as ApoBD and Apoptotic EV(44–46). In addition, there is growing evidence that the formation and secretion of calcifying vesicles are critical steps for vascular mineralization(47–50). We next examined whether IKK2 deficiency induces the secretion of calcifying mediators such as ApoBD. Treatment of wild-type VSMCs with a 48 hour condition media from IKK2KO VSMC cultures significantly induced levels of matrix calcium contents by 3.8-fold compared to treatment with media from wild-type VSMC cultures (Figure 7A), suggesting that IKK2 deficiency induces the secretion of calcifying factors. Since ApoBD has known to induce mineralization of VSMCs(40), the condition media from wild-type and IKK2KO VSMCs was fractionated by ultracentrifuge. Interestingly, treatment of wild-type VSMCs with ApoBD-enriched fractions from either wild-type or IKK2KO cultures did not affect mineralization of VSMCs, whereas EV enriched fractions from IKK2KO VSMC cultures but not wild-type VSMC cultures significantly induced the mineralization of VSMCs (Figure 7B). NTA analysis revealed that IKK2 deficiency enhanced the secretion of EV by about 5-fold under normal and TNFα-treated conditions (Figure 7C). IKK2 deficiency did not affect the size distribution of EV compared to wild-type VSMCs (Figure 7D and 7E). To confirm that IKK2 deficiency increases the secretion of EV, levels of EV markers in the culture media were determined by immunoblot analysis. Consistent with NTA, IKK2 deficiency increased levels of CD63, CD9, annexin A2 and annexin 6 in the cultured media by 185-fold, 697-fold, 57-fold and 12-fold, respectively (Figure 7F and Supplemental Figure 4A-4D). We next examined whether cell death is linked to IKK2KO-mediated mineralization, osteogenic differentiation and EV secretion. Recent studies demonstrated that treatment with the cell death inhibitor GSK2656157 completely blocked TNFα-induced cell death. As shown in Figure 7G/7H and Supplemental Figure 4E/4F, treatment with the cell death inhibitor totally blocked TNFα- and high-phosphate-induced mineralization and osteogenic differentiation of both wild-type and IKK2KO mice. Blocking cell death significantly reduced the secretion of EV from wild-type and IKK2KO cells (Figure 7I). In addition, IKK2 deficiency increased levels of ALP protein in CD63-positive EVs, but that was significantly reduced by blocking cell death (Figure 7J and 7K), suggesting that calcifying EVs were increased by IKK2KO-mediated cell death. We also tested whether IKK2-NFκB modulations affects levels of calcifying EVs in the aortic media in vivo. Immunofluorescence analysis of the aortic sinuses showed that the number of CD63^+^ALP^+^ double positive puncta were increased in the aortic media of CKD SMC-IKK2KO mice, and reduced by SMC-IκBα deficiency (Figure 7L and 7M).

**Figure 6.**
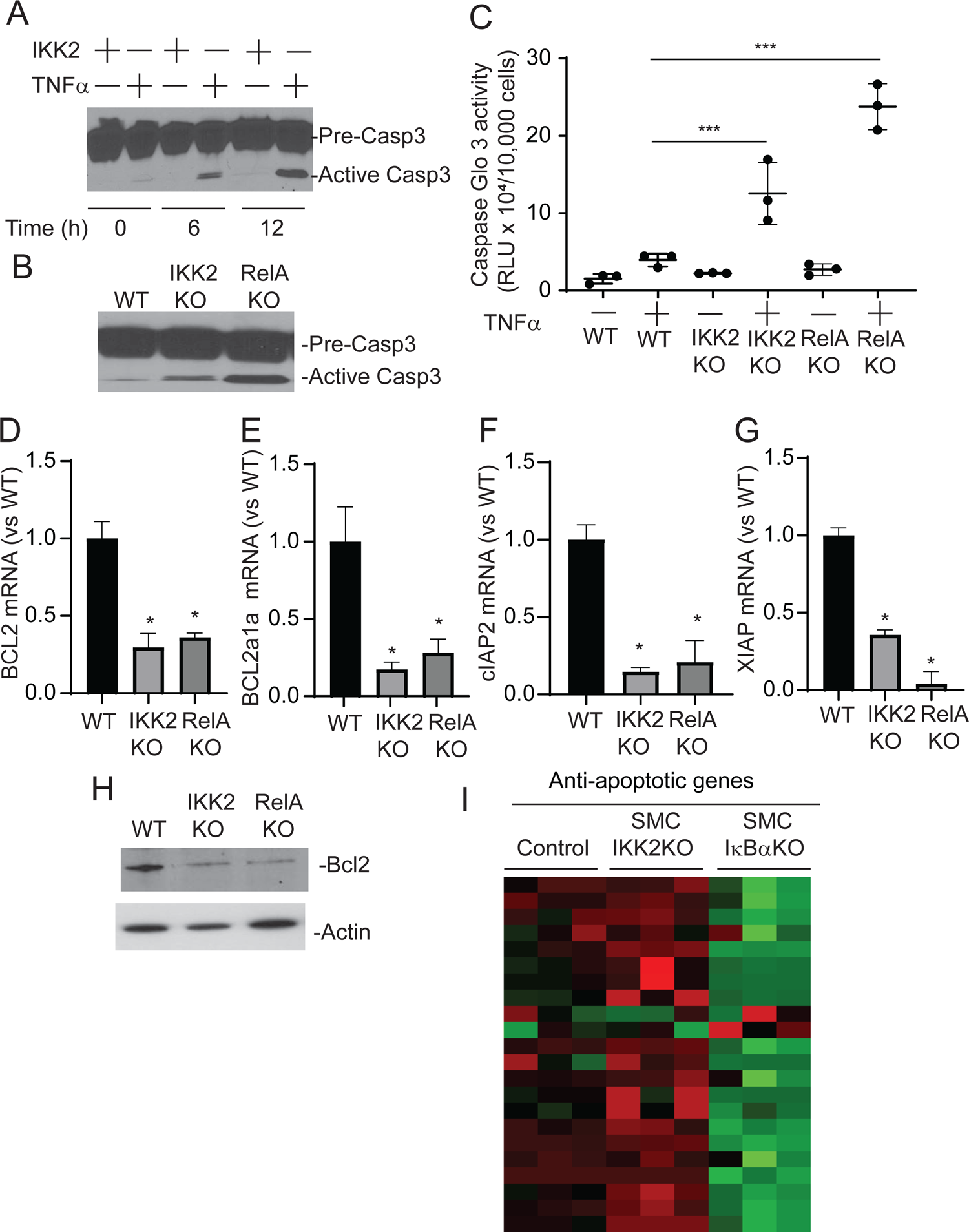
Deletion of the IKK2-NFκB pathway induces apoptosis of VSMCs. A) Immunoblot analysis of Caspase 3 (Casp3) in IKK2KO VSMCs treated with TNFα. VMSCs were treated with TNFα for 6 hours and 12 hours. B) Immunoblot analysis of Casp3 in IKK2KO and RelAKO VSMCs treated with high-phosphate. VMSCs were treated with high-phosphate for 12 hours. C) Casp3 activity in IKK2KO and RelAKO VSMCs treated with TNFα. VMSCs were treated with TNFα for 12 hours. D-G), mRNA levels of anti-apoptotic proteins in IKK2KO and RelAKO VSMCs. H) Immunoblot analysis of Casp3 in IKK2KO and RelAKO VSMCs. VMSCs were treated with TNFα for 12 hours. I) Heatmap with mRNA levels of 21 anti-apoptotic proteins in the aortic media of SMC-IKK2KO and SMC-IκBKO mice. Aortic medial layers were dissected 3 weeks after CKD was induced. The numbers are shown in Supplemental Table 4. *P<0.05 and ***P<0.001

**Figure 7.**
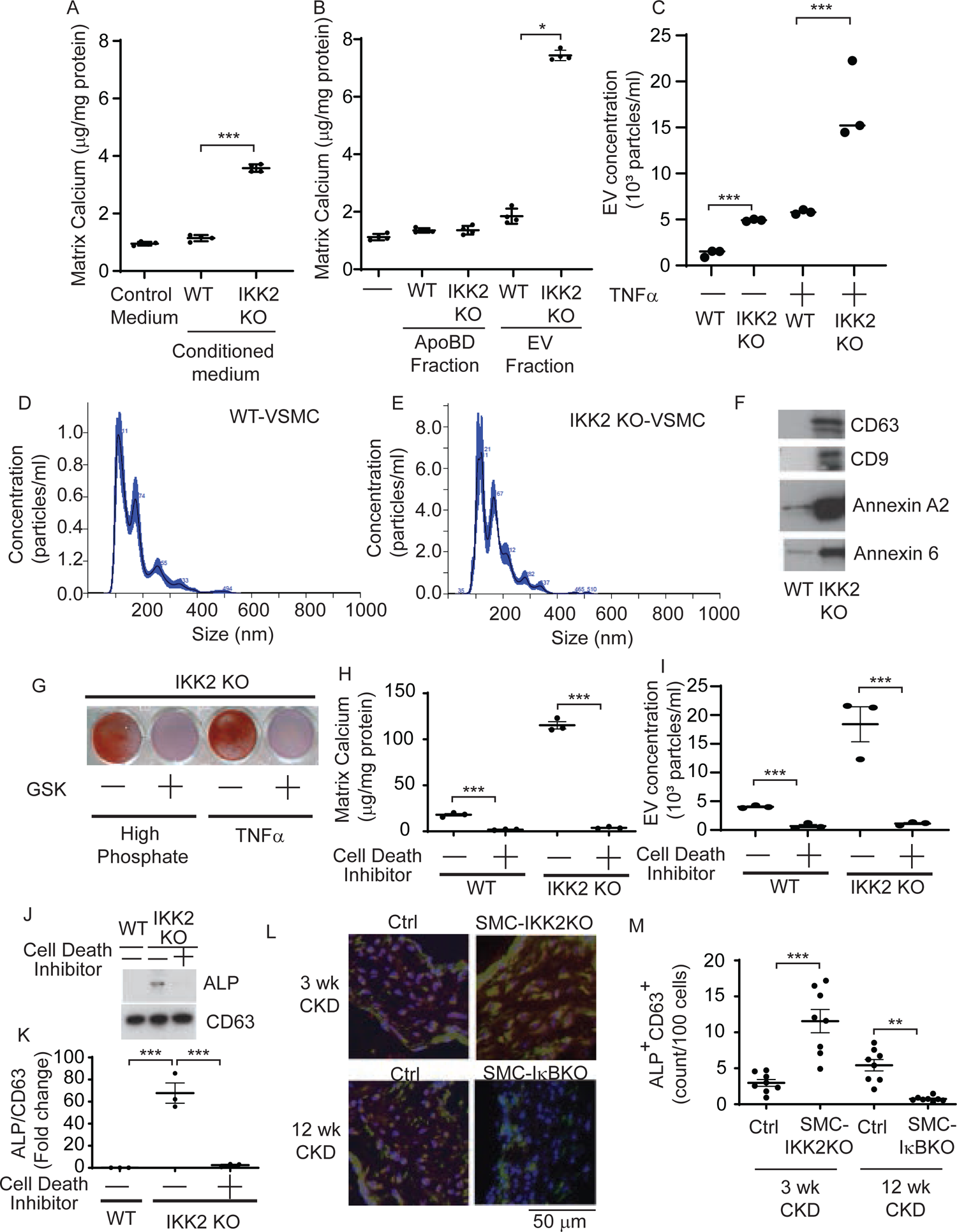
Deletion of IKK2 induces the secretion of claying extracellular vesicles that is completely inhibited by cell death inhibitor. A) Condition media from IKK2KO VSMCs but not wild-type VSMCs induced mineralization of wild-type VSMCs. The conditioned media were collected from IKK2KO and wild-type VSMCs treated with TNFα for 48 hours in the presence of EV-depleted FBS. Wild-type VSMCs were treated with the conditioned media for 7 days. B) EV-enriched fraction but not ApoBD fraction induces mineralization of VSMCs. Wild-type VSMCs were treated with 40μg/ml ApoBD- and EV-enriched fractions from IKK2KO and wild-type VSMC cultures for 7 days. ApoBD-enriched and EV-enriched fractions were isolated with sequential centrifugation. C) EV concentration and D, E) size in IKK2KO VSMCs. EV concentration was determined with NTA using a Nano-Sight NS500. VSMCs were treated with TNFα for 48 hours. F) Immunoblot analysis of EV markers (CD63, CD9, Annexin A2 and Annexin 6) in the culture media of IKK2KO VSMCs. VSMCs were treated with TNFα for 48 hours. Culture media (20 μl) were subjected to immunoblot analysis. The densitometry quantitations are shown in Supplemental Figure 4. G) Alizarin Red stain, H) matrix calcium content and I) EV concentration in IKK2KO VSMCs treated with cell death inhibitor. VSMCs were treated with high phosphate or TNFα in the absence and presence of cell death inhibitor (0.1 μM, GSK2656157) for 7 days for calcium analysis and 2 days for EV analysis. J) Immunoblot analysis and K) quantitation of ALP in the EV fraction of IKK2KO VSMCs treated with cell death inhibitor for 48 hours. L, M) Immunofluorescence analysis of aortic calcifying EV in SMC-IKK2KO and SMC-IκBKO mice. CD63 (Green) and ALP (Red) double positive puncta (Yellow, CD63^+^ALP^+^) were analyzed as calcifying EV. *P<0.05 and ***P<0.001

## Discussion

IKK2-NFκB signaling is activated by numerous discrete stimuli and is a master regulator of the inflammatory response(17–19). Activation of the IKK2-NFκB pathway has been proposed to contribute to the etiology of cardiovascular complications in CKD such as vascular calcification(24–30). However, the VSMC-specific role of IKK2-NFκB in the regulation of CKD-induced vascular complications is still obscure. In this study, mRNA-seq analysis coupled with cell sorting, immunoblot analysis, and EMSA analysis revealed that CKD induces >100 genes involved in the inflammatory response locally in VSMCs via activation of the IKK2-NFκB pathway. Pro-inflammatory genes increased by CKD were associated with medial calcification and vascular stiffness. Based on previous studies from other groups, we expected that inhibition of the NFκB pathway attenuates vascular mineralization by reducing the production of pro-inflammatory mediators locally from VSMCs. In fact, all of the in vitro and in vivo models tested in this study suggest that activation of IKK2-NFkB signaling by CKD induces inflammatory mediator expression in VSMCs. Unexpectedly, however, we demonstrated that activation of the IKK2-NFκB pathway in VSMCs elicits strong protective effects for CKD-dependent vascular complications. Using the CRISPR-Cas9 technique, we first deleted 3 key molecules (IKK2, RelA and NFKB1) involved in IKK2-NFκB signaling from cultured VSMCs. All of the gene deletions aggravated TNFα- and high Pi-induced mineralization and osteogenic differentiation of VSMCs. We used two mouse models to modulate the IKK2-NFkB pathway specifically in VSMCs. SMC-specific IKK2KO mice had significantly reduced CKD-induced NFκB activation and pro-inflammatory mediator expressions in VSMCs; SMC-specific IKK2 deficiency accelerated CKD-induced medial calcification. On the other hand, SMC-IκBKO deficiency attenuated CKD-mediated vascular complications despite the activation of NFκB and pro-inflammatory mediator expressions in VSMCs. These findings strongly suggest that the activation of IKK2-NFκB signaling in VSMCs is a defense mechanism against calcified vascular stiffness in CKD.

We provided mechanistic insights into the protective role of IKK2-NFκB signaling in regulating CKD-mediated vascular calcification. A number of recent studies have proposed that the secretion of calcifying extracellular vesicles containing CD63, annexins and ALP is a pivotal event in the pathogenesis of vascular osteogenesis and mineralization(48–50). Our studies reveal that the inhibition of IKK2 increased the secretion of calcifying EV. Mechanistically, cell death by the inhibition of the IKK2-NFκB pathway activates the secretion of calcifying EVs from VSMCs, resulting in vascular mineralization and osteogenesis. It was previously considered that the secretion of apoBD mediates cell death-induced VSMC mineralization. Since ApoBD fractions of IKK2KO cells did not influence the mineralization of VSMCs, apoBD plays a minor role in the IKK2-NFκB-cell death-mineralization cascade. In this study, we used GSK2656157 as a cell death inhibitor because the chemical completely and potently (∼nM level) blocks TNFα-mediated cell death by inhibiting multiple protein kinases such as PERK and RIPK1(57). We have previously shown that PERK-mediated ER stress is involved in TNFα-mediated vascular calcification(14). Interestingly, cell death inhibitor treatment blocked TNFα-but also high-phosphate-mediated VSMC mineralization. In addition, GSK2656157 inhibited other uremic toxin (Cresol sulfate and Indoxyl sulfate)-induced mineralization (data not shown). Taken together, these data suggest that VSMC cell death is a critical step in the pathogenesis of CKD-dependent medial calcification. The mechanism by which deletion of the IKK2-NFκB pathway induces apoptosis is due to the reduction of anti-apoptotic genes. We have found in this study that the activation of NFκB by the deletion of IκB significantly induces 21 out of 22 anti-apoptotic genes expressed in VSMCs.

CKD induces chronic systemic inflammation and activates the NFκB pathway in other tissues and cells such as kidney, lung and adipose tissues in addition to VSMCs(51, 58). We and other groups have previously shown that blocking specific pro-inflammatory cytokines such as TNFα is effective in preventing vascular calcification in animal models(14, 15). We still believe that the inhibition of systemic inflammation is beneficial for blocking CKD-mediated medial calcification. Several previous studies have also shown that systemic inhibitions of the IKK2-NFκB pathway by chemical inhibitors or RNAi also block vascular calcification(28, 55). Monocytes could be a major source of increased pro-inflammatory cytokines in CKD, although unlike atherosclerotic CKD models, macrophage infiltrations in the aorta were not observed in our CKD mouse model of medial calcification. Further studies are required for identifying how tissue/cell inflammation and pro-inflammatory production play a major role in regulating CKD-induced medial calcification.

In this study, we unexpectedly demonstrate that activation of the IKK2-NFkB pathway works as a safeguard for VSMCs from ectopic mineralization in CKD by blocking cell death-mediated activation of calcifying extracellular vesicle secretion. In addition, this study suggests that pro-inflammatory cytokines produced locally from VSMCs have more minor effects on the pathogenesis of CKD-mediated medial calcification than we previously anticipated.

## METHODS

### Animals

SMMHC-GFP (stock# 7742), SMMHC-CreER^(T2)^ (stock# 19079) mice and CKD mice were generated as previously described. IKK2 conditional knockout (IKK2^F/F^) mice were kindly provided by Dr. Karin at the University of California San Diego(59). To generate IκBα (nuclear factor of kappa light polypeptide gene enhancer in B-cells inhibitor, alpha) conditional KO mice, we flanked exons 2-5 of the NFKBIA gene with two *lox*P sites (Supplemental Figure 3). The targeting vector PG00171_Y_4_H09–Nfkbia was purchased from MMRRC. The targeting vector was linearized with AsiSi and introduced by electroporation into murine B6/129 hybrid EC7.1 embryonic stem cells. Karyotypically normal ES clones were microinjected into C57BL/6 blastocysts to produce chimeric founders at the Bioengineering Core Facility at the University of Colorado-Denver. The IkBα^F/F^ mice were crossed with Rosa-Flp mice to remove the LacZ-Neo cassette. All of the mouse strains were backcrossed at least 10 times with DBA/2J mice that are susceptible to CKD-dependent medial calcification. The DBA genetic background was checked with the PCR speed congenic service at DERC Molecular Biology Core Facility on our campus. To generate VSMC-specific IKK2 and IκBα KO mice, IKK2^F/F^ and IκBα^F/F^ mice were intercrossed with SMMHC-CreER^(T2)^ mice to obtain SMMHC-CreER^(T2)^; IKK2^F/F^ mice and SMMHC-CreER^(T2)^; IκBα^F/F^ mice, respectively. SMMHC-CreER^(T2)^ mice were used as control mice. One week after the surgeries, CKD mice were injected intraperitoneally with tamoxifen (40 mg/kg body weight) in vegetable oil. After the injections, VSMC-specific IKK2 KO (SMC-IKK2KO) and VSMC-specific IκBα KO (SMC-IκBαKO) mice were maintained on a special diet (TD110198) for 3 and 12 weeks, respectively, unless we indicated otherwise. As previously described, calcified lesions in the aortic arches were analyzed as previously described using von Kossa staining. Cell death was detected using an In Situ Cell Death Detection Kit (Roche, catalog # 12156792910). CD63 and ALP double positive (CD63^+^; ALP^+^) calcifying extracellular vesicles in the aortic sinus were detected using Alexa 488-conjugated CD63 polyclonal antibody (Novus, MX-49.129.5) and Alexa 568-conjugated ALP recombinant antibody (Selleck, A511) as previously described(39, 42, 54, 60). At least 5 sections and 100 DAPI-positive nuclei from each sample were analyzed.

### Aortic pulse wave velocity (aPWV)

aPWV was assessed non-invasively using an Indus Doppler Flow Velocity System (Scintica, Canada) as previously described(61, 62). Briefly, isoflurane (2%) was used to anesthetize mice that were placed supine with legs secured to ECG electrodes on a heated board. Doppler probes were placed on the skin at the transverse aortic arch and abdominal aorta ∼ 4 cm apart. For each site, the pre-ejection time, or time between the R-wave of the ECG to the foot of the Doppler signal was determined. To calculate aPWV, the distance between the probes was divided by the difference in the thoracic and abdominal pre-ejection times and is presented as centimeters/s (cm/s). Following aPWV measures, mice were euthanized by exsanguination via cardiac puncture while anesthetized with isoflurane.

### Cell cultures

Human VSMCs and mouse VSMCs were purchased from Applied Biological Materials (T0515, Richmond, Canada) and American Type Culture Collection (ATCC), respectively. VSMCs were maintained in DMEM containing 1% Exosome-depleted FBS (A2720801, ThermoFisher) with 100 U/ml penicillin and 100 μg/ml streptomycin. VSMCs were treated with either 1 ng/ml human TNFα (GenScript, Z00100) or 2.4 mM inorganic phosphate in the absence or presence of 0.1μM GSK2656157 (Cayman, 17372).

### CRISPR-Cas9 system-based gene knockout of IKK2, IκBα and NFKB1 genes

IKK2, IκBα and NFKB1 gene sgRNA was cloned into LentiCRISPRv2 plasmid (Addgene) as previously described(54, 60). The sgRNA sequences are shown in Supplemental Table 5. HEK293T cells were seeded at 6 x 10^5^ cells/well in 6-well plates, grown overnight, and then transfected with 300 ng psPAX2, 100 ng pMD2 and 400 ng of each sgRNA CRISPR Cas9 lentivirus plasmid (plasmid amount rate 3:1:4) using Turbofect transfection reagent (Thermo Fisher Scientific). Lentiviral media was centrifuged once at 1500 x g for 3 minutes and the supernatant was collected. VSMCs were seeded in 6-well plates and infected 24 hours later with each lentiviral media in the presence of 10 μg/ml polybrene. Cells were treated with 5 μg/ml puromycin for selection of infected cells. Total RNA of heterogeneous cells was collected and cDNA synthesis was conducted from the RNA template, followed by high resolution melting analysis with a StepOne Plus qPCR instrument (Applied Biosystems) to check for mutations occurring on regions around IKK2, IκBα and NFKB1 sgRNA target sequences. Heterogeneous cells with gene mutations were plated at 0.5 cells/well in a 96-well plate to obtain a single cell clone. Protein from gene edited clones was prepared and analyzed by immunoblot analysis to determine whether gene knockout was complete.

### Generation of VSMCs expressing IKK2DN

FLAG-IKK2 kinase inactive K44M (IKK2DN, Addgene, #15466) was cloned into pLenti-CMV-Puro DEST vector (Addgene, #17452) using a Gateway cloning system (Invitrogen). VSMCs were infected with lentivirus containing FLAG-IKK2DN. Cells were treated with 5 μg/ml puromycin for selection of infected cells. The single clones were analyzed by immunoblot analysis with FLAG antibody (M2, Sigma-Aldrich).

### RNA analysis

Total RNA was isolated using a Direct-zol RNA kit (Zymo Research). cDNA was synthesized from 500 ng total RNA using a High-Capacity cDNA Reverse Transcription Kit (Applied Biological Materials Inc). qRT-PCR was performed using an Applied Biosystems StepOne Plus qPCR instrument with SYBR™ Select Master Mix according to the manufacturer’s instructions. Primer sequences are shown in Supplemental Table 5.

### Calcium content in cultured cells and aortas

For evaluation of vascular mineralization, VSMCs were plated at 1.0 x 10^5^ cells/well in a 12-well plate and grown overnight. VSMCs were treated with 1 ng/ml TNFα or 2.4 mM Pi every 2 days for 6 days. VSMCs were incubated with 0.6 N HCl overnight at 4°C. After incubation, 0.6 N HCl was collected to measure calcium content, and then VSMCs were lysed with 0.1 N NaOH/0.1% SDS to measure protein concentration with a BCA assay. Aortas were collected from mice and stored at −20°C. Dried aorta was defatted with chloroform and methanol (2:1) for 48 hours and dehydrated with acetone for three hours. The dried samples were incinerated to ashes at 600°C for 24 hours using an electric muffle furnace (Thermo Scientific), then extracted with 0.6 N HCl. Calcium content from cultured cells and aortas was quantified using the o-cresolphthalein method. In addition, VSMCs were stained with Alizarin Red to identify calcium deposits 6 days after TNFα and Pi treatments(42, 53, 54).

### RNA-seq

Total RNA was isolated using a Direct-zol kit. mRNA-seq library construction and sequencing was performed at BGI America (http://www.bgi.com) in accordance with the manufacturer’s instructions using a DNBseq system as previously described(60, 63). The raw data were deposited to the NIH Gene Expression Omnibus as GSE229679.

### Cell sorting

To isolate aortic VSMCs, isolated aortas were digested to single cells by digestion at 37°C in collagenase buffer (3.2 mg/ml collagenase II, 0.7 mg/ml elastase, 0.2 mg/ml soybean trypsin inhibitor) in Hank’s buffered saline solution as previously described(42, 53). Aortas were harvested under sterile conditions following flushing of the vasculature system with sterile heparinized PBS and minced prior to digestion. Single cell suspension was sorted based on GFP for SMMHC-GFP mice under sham-operation and CKD. Sorting was performed on a MoFlo high-speed cell sorter at the University of Colorado flow cytometry and sorting core facility.

### Electrophoretic mobility shift assay

Electrophoretic mobility shift assay (EMSA) analysis was performed as previously described. The DNA binding activity of NF-κB was assayed according to the protocol from Promega Corp. Briefly, the oligo with NF-κB consensus binding element (Promega) was end-labeled by T4 polynucleotide kinase (Promega) using [γP^32^]-ATP (BioRad). Thirty μg of total protein extract was isolated from the VSMCs using 1X passive lysis buffer (Promega) and was mixed with radio-labeled oligo for binding. Unlabeled cold probe was used to compete with the radio-labeled probe to show binding specificity. The reaction mixture was loaded to a 5% polyacrylamide gel under non-denaturing conditions and was separated by electrophoresis at 4°C. The gel was then dried and exposed to X-ray film to visualize the binding of NF-κΒ onto the radio-labeled probe. The binding specificity was previously shown by blockage of binding with excessive competitive cold probe, and the position of the NF-κB p65/50 complex was confirmed using anti-p65 (#8242, Cell Signaling Technology) and anti-p50 antibodies (14-6732-81, Thermo Fisher Scientific)(64, 65).

### Immunoblot analysis

Cell and tissue lysates were prepared using RIPA buffer (150 mM NaCl, 1% Nonidet P-40, 0.5% sodium deoxycholate, 0.1% SDS, 50 mM Tris, pH 8.0). Cells were disrupted by pipetting 10-15 times, centrifuged at 13,800 x g for 10 minutes at 4°C and the supernatant was collected for total cell lysates. The samples were separated by SDS-PAGE, transferred to a nitrocellulose membrane, and immunoblotted with the following antibodies: IKK2 (D30C6, #8943), p-IKK2 (16A6, #2697), IκB (44D4, #4812), p-IκB (14D4, #2859), p65 (D14E12, #8242), p-p65 (93H1, #3033), NFKB1 (D4P4D, #13586), Annexin A2 (D11G2, #8235) and Casp3 (#9662) from Cell Signaling Technology, Gapdh (V-18) from Santa Cruz Biotechnology, and β-Actin (66009) from Proteintech. Annexin 6 (A305-309A), CD63 (NVG-2), and TNAP (A511) antibodies were purchased from Bethyl, Biolegend and Selleck, respectively. Samples were visualized using horseradish peroxidase coupled to appropriate secondary antibodies with enhancement by an ECL detection kit (Thermo Fisher Scientific).

### Nanoparticle tracking analysis (NTA)

NTA was performed at the University of North Carolina Nanomedicines Characterization Core Facility in accordance with the manufacturer’s instructions using NanoSight NS500 (Malvern Panalytical).

### Isolation of apoptotic bodies and extracellular vesicle fraction

A 200 ml cell culture medium was centrifuged at 300 x g for 10 minutes twice. The collected supernatant was centrifuged for 20 minutes at 3,000 x g twice. The pellet was resuspended in 1x PBS as the Apoptotic bodies (ApoBD)-enriched fraction. The 3,000 x g supernatant was centrifuged for 20 minutes at 15,000 x g twice. The collected supernatant was further ultracentrifuged for 1 hour at 100,000 x g. The 100,000 x g pellet was washed with 1x PBS and re-centrifuged for 1 hour at 100,000 x g as the extracellular vesicles (EV)-enriched fraction(48, 66, 67).

### Statistical analysis

Data were collected from more than two independent experiments and reported as the means ± S.E.M. Statistical analysis for two-group comparison was performed using the Student’s *t* test, or one-way ANOVA or two-way ANOVA with a Newman-Keuls post-hoc test for multi-group comparison. Significance was accepted at *P* < 0.05.

### Study Approval

All animal protocols and experimental procedures were approved by the Institutional Committees at the University of Colorado-AMC.

### AUTHOR CONTRIBUTIONS

Conceptualization, M. Miyazaki; Methodology, M. Masuda, S.M-A., Y.S. and M. Miyazaki; Formal Analysis, M. Masuda, S.M-A, and M. Miyazaki; Data Curation, M. Masuda, S.M-A, and M. Miyazaki; Investigation, M. Masuda, S.M-A A.L.K, Y.S., M. Miyazaki.; Writing – Original Draft, M. Miyazaki; Writing – Review & Editing, A.L.K., Y.S., M. Masuda and M. Miyazaki; Visualization, M. Masuda, S. M-A., M. Miyazaki; Funding Acquisition, M. Miyazaki; Supervision, M. Miyazaki.

## ACKNOWLWGEMENTS

The authors’ work was supported by grants from NIH HL132318, R01HL1157069 and R01DK124901 to M. Miyazaki.

