## Supplemental Figures for "Activation of the IKK2-NFκB pathway in VSMCs inhibits calcified vascular stiffness in CKD by reducing the secretion of calcifying extracellular vesicles"

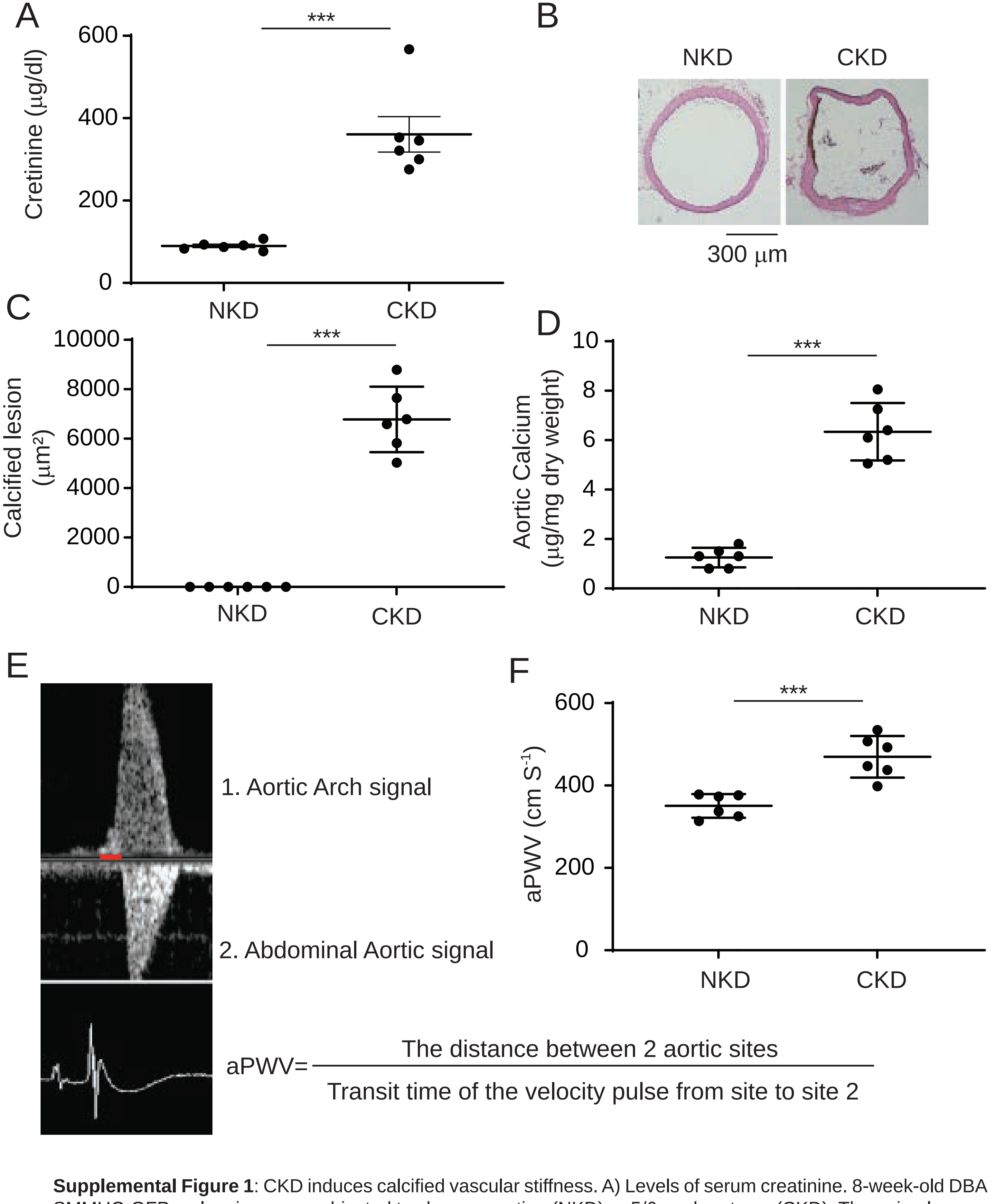


**Supplemental Figure 1**: CKD induces calcified vascular stiffness. A) Levels of serum creatinine. 8-week-old DBA SMMHC-GFP male mice were subjected to sham operation (NKD) or 5/6 nephrectomy (CKD). The animals were euthanized 12 weeks after the surgeries. Levels of serum creatinine were analyzed with LC-MS/MS. B) Histological analysis of aortic arches with von Kossa staining. C) The quantitation of calcified lesions in CKD mice. D) Aortic calcium content was analyzed with ash assay coupled with calcium colorimetric assay. E, F) aPWV. aPWV was analyzed using an Indus Doppler Flow Velocity System 12 weeks after the surgeries. ***P<0.001

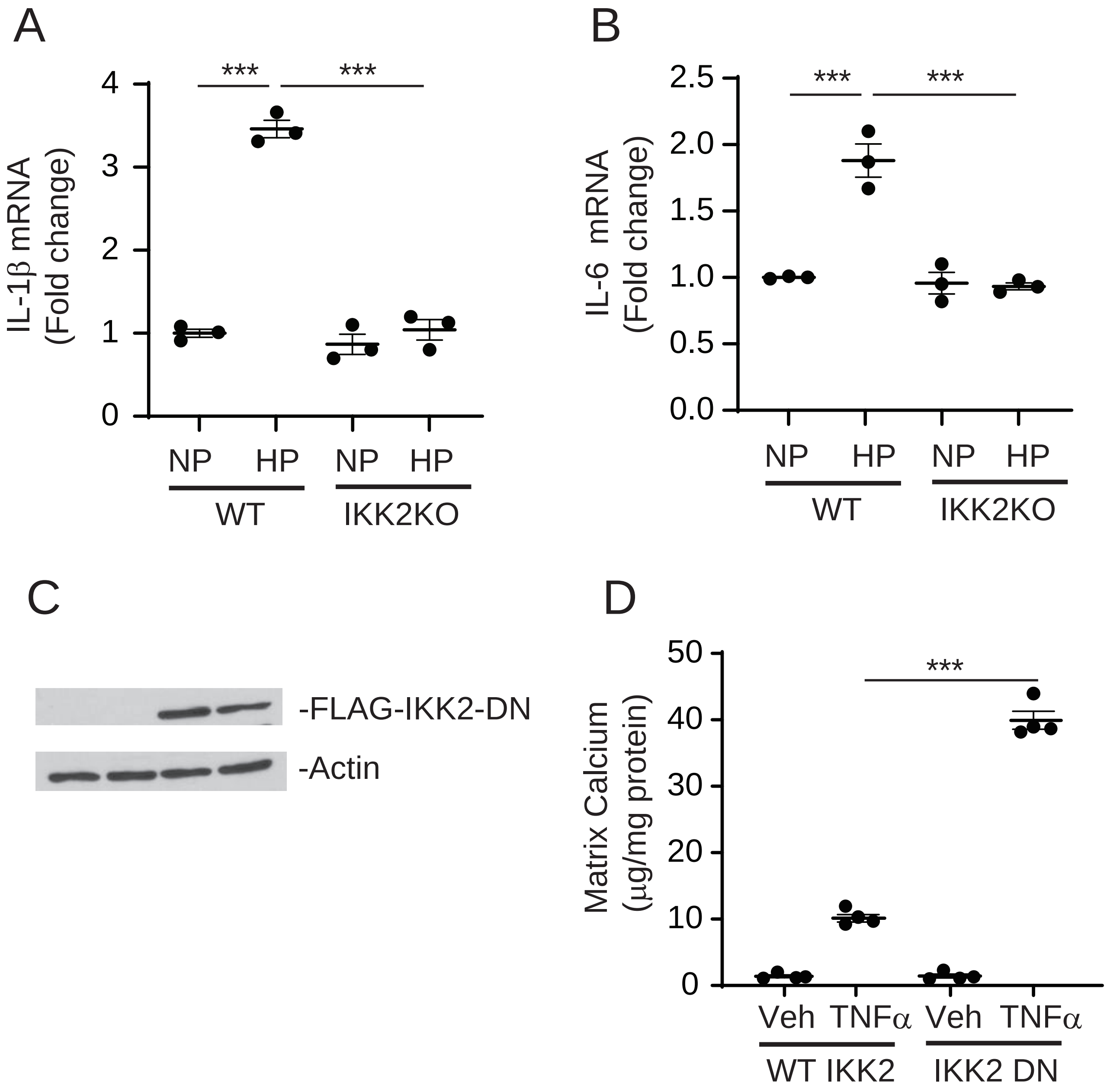


**Supplemental Figure 2:** IKK2 knockout and IKK2 dominant negative expression induces vascular calcification. A, B) mRNA levels of IL-1β and IL-6 in IKK2KO VSMCs treated with high-phosphate. VSMCs were treated with high-phosphate (2.4mM) for 16 hours. C) Immunoblot analysis of FLAG-IKK2DN. D) Levels of matrix calcium. VSMCs were treated with TNFα for 6 days. ***P<0.001

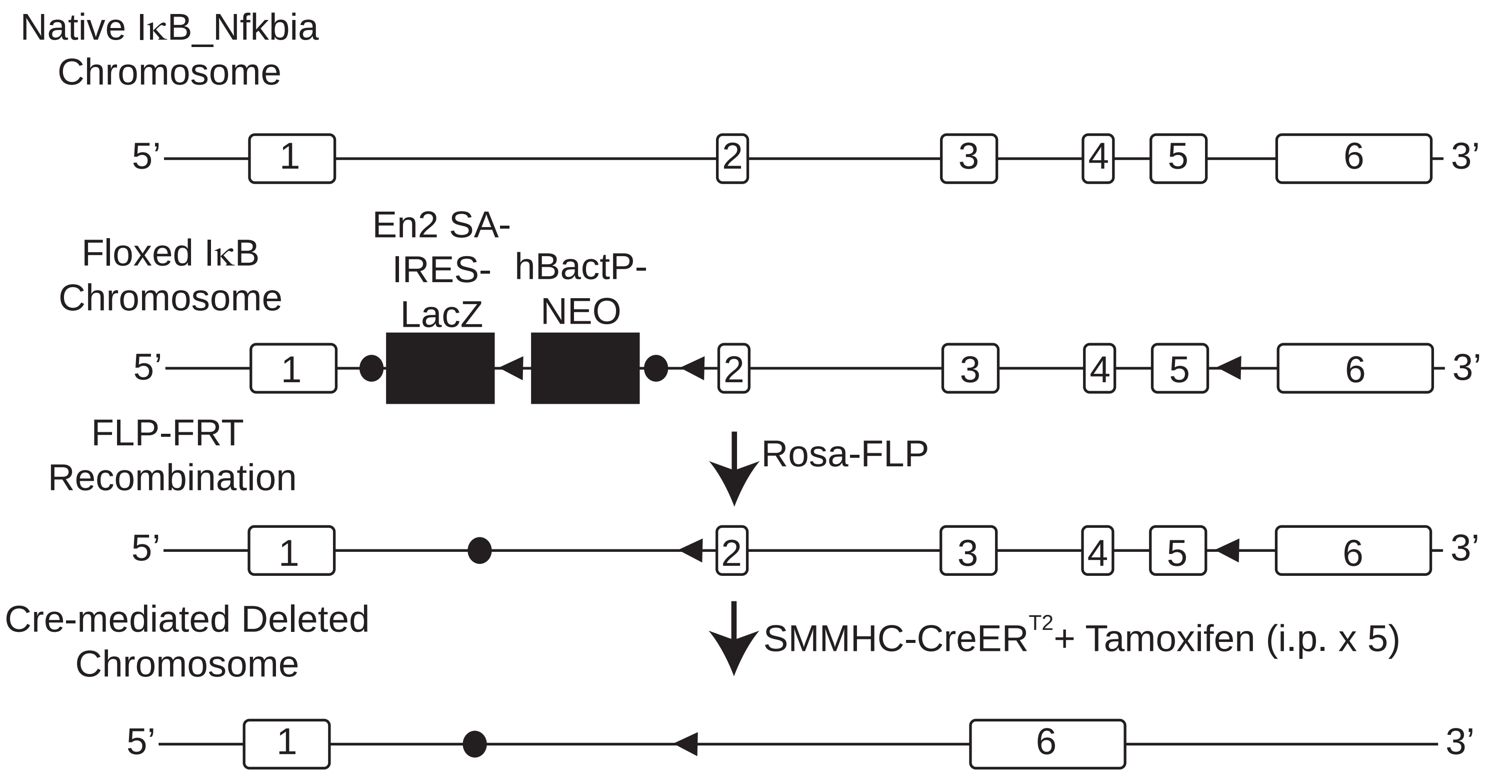


**Supplemental Figure 3:** The strategy of generation of SMC-IκBKO mice.

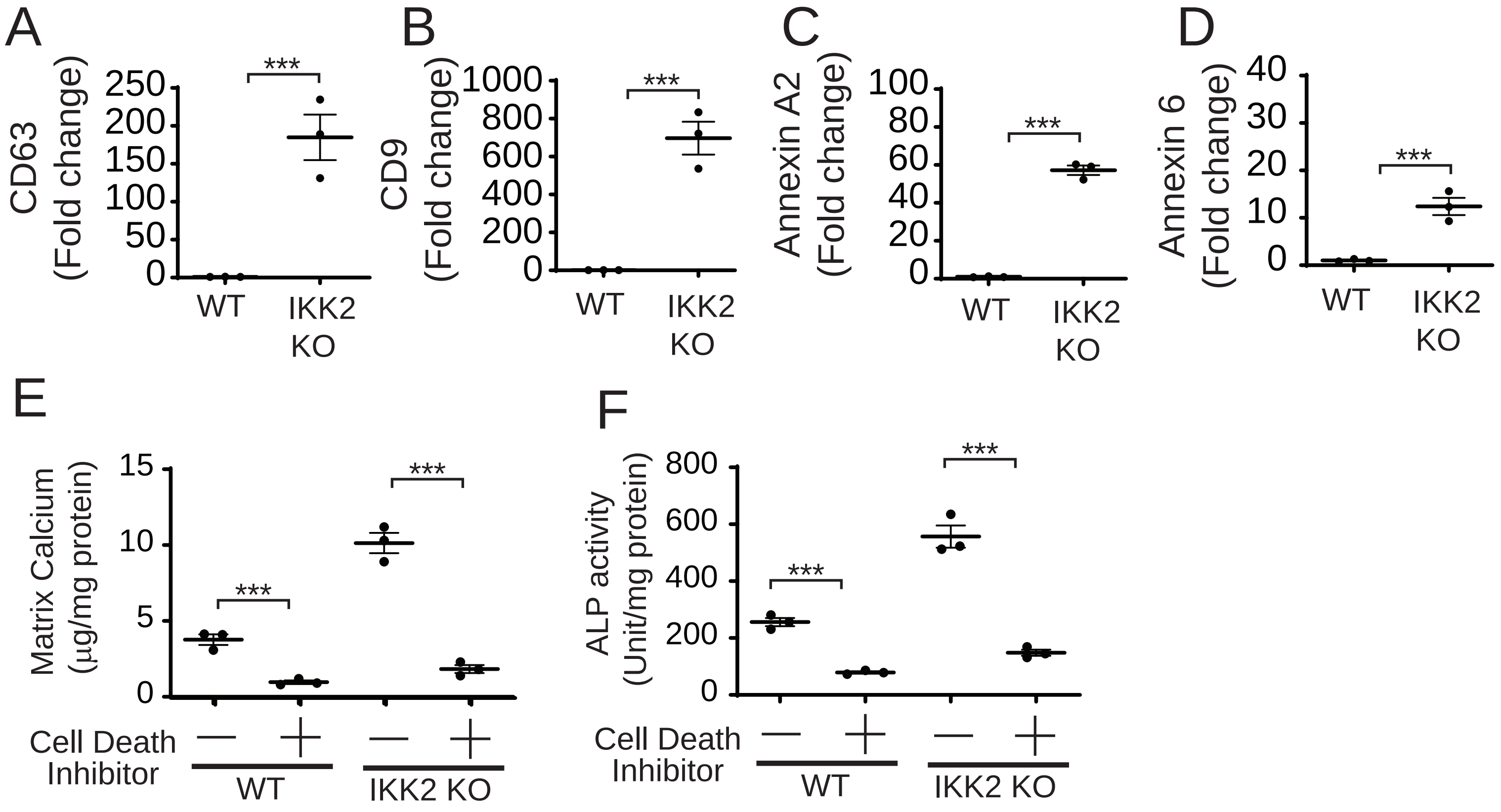


**Supplemental Figure 4:** Inhibition of cell death blocks IKK2 deficiency-induced mineralization and EV secretion in VSMCs. A-D) Quantitation of EV markers (CD63, CD9, Annexin A2 and Annexin 6) in the culture media of IKK2KO VSMCs. VSMCs were treated with TNFα for 48 hours. Culture media (20 μl) were subjected to immunoblot analysis. E) Matrix calcium content in IKK2KO VSMCs treated with cell death inhibitor. VSMCs were treated with 2.4 mM high phosphate in the presence of cell death inhibitor (0.1 μM GSK2656157) for 7 days. F) ALP activity in IKK2KO VSMCs treated with cell death inhibitor. VSMCs were treated with 1 ng/ml TNF in the presence of a cell death inhibitor (0.1 μM GSK2656157) for 7 days. ***$P<0.001$.

Fig 1C

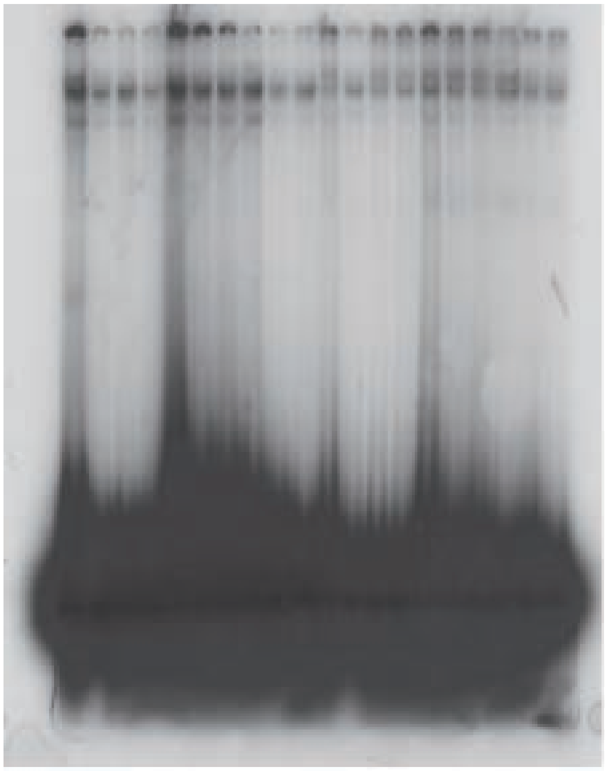

Fig 1D

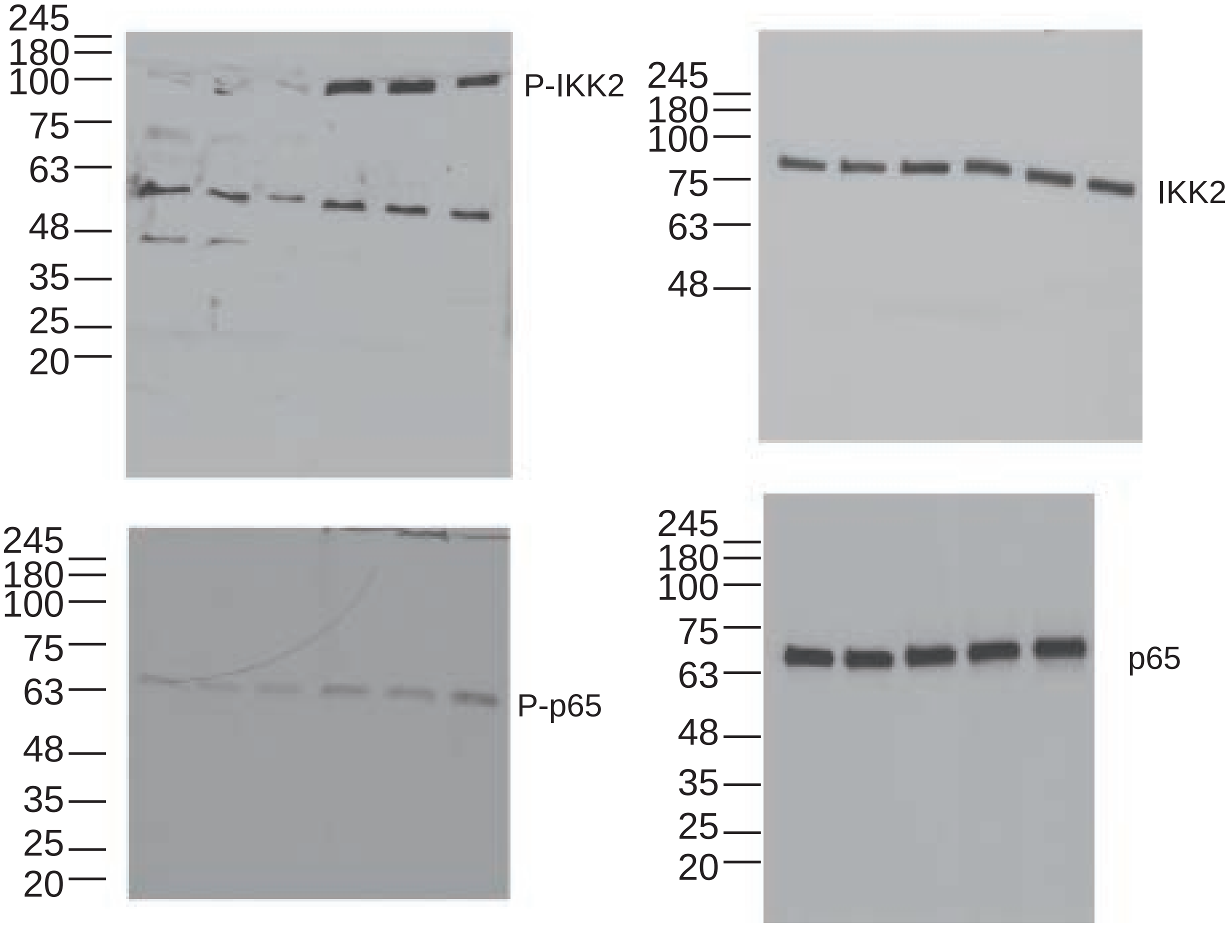
245
180
100
75
63
48
35
25
20
P-IKK2
245
180
100
75
63
48
IKK2
245
180
100
75
63
48
35
25
20
P-p65
245
180
100
75
63
48
35
25
20
p65

Fig 3A

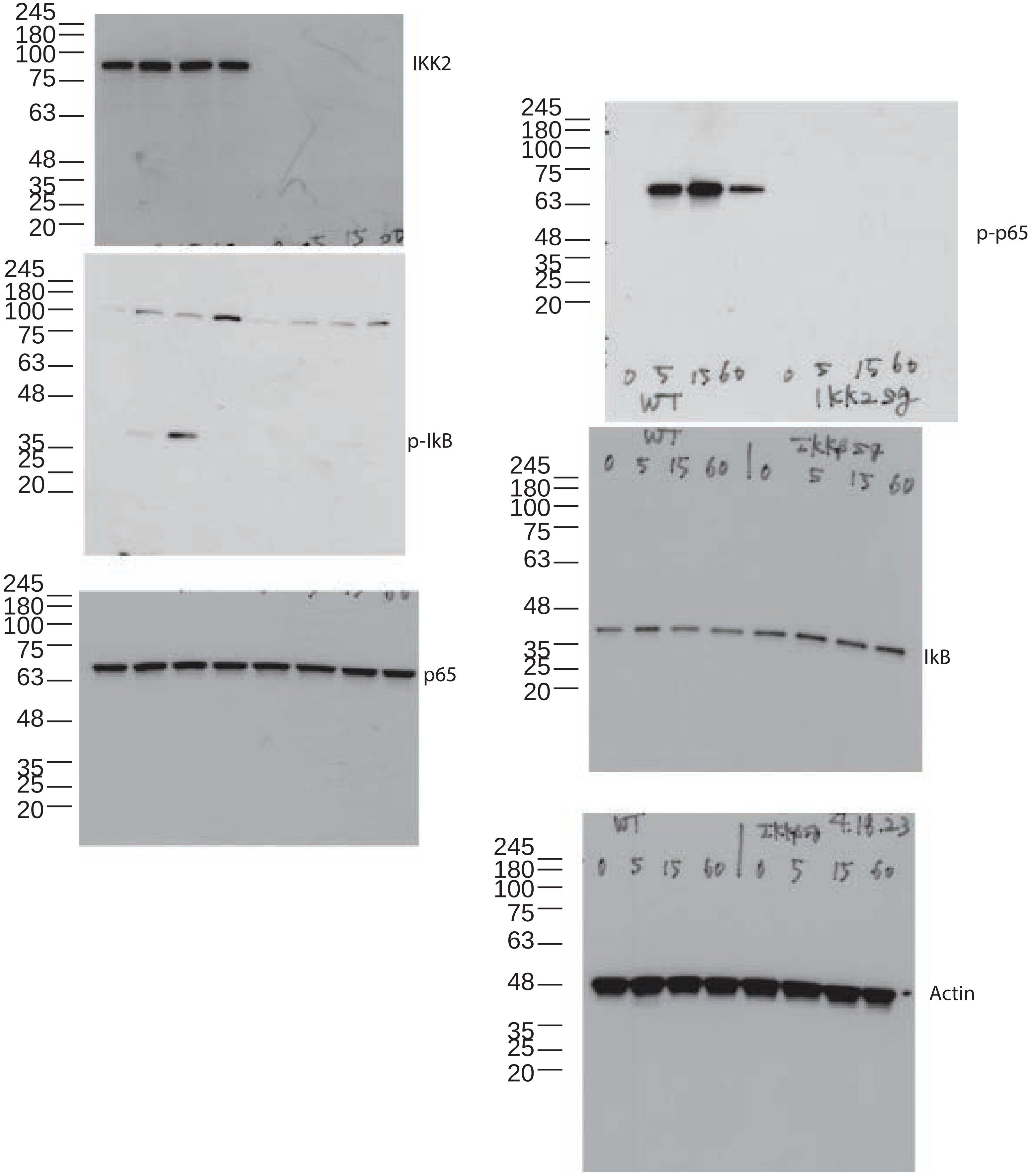

245
180
100
75
63
48
35
25
20
IKK2
p-IkB
p65
p-p65
IkB
Actin
0 5 15 60
WT
0 5 15 60
Ikk2sg
WT
0 5 15 60
0 5 15 60
4.18.23

S.Fig 2

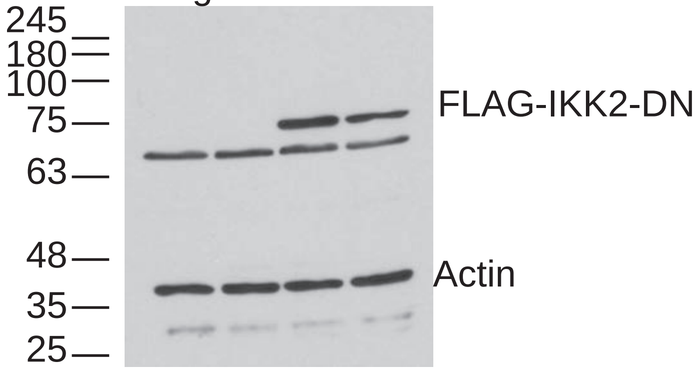


Fig 3A

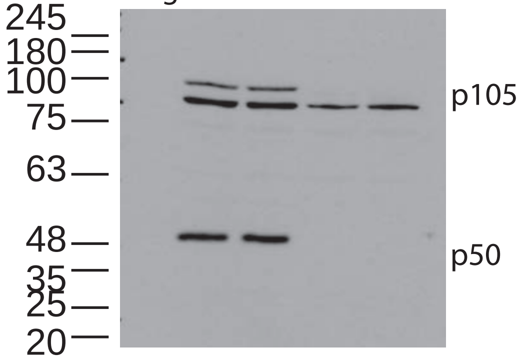


Fig 3B

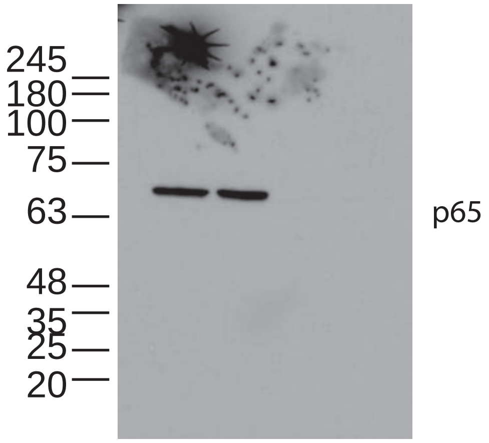


Fig 3A

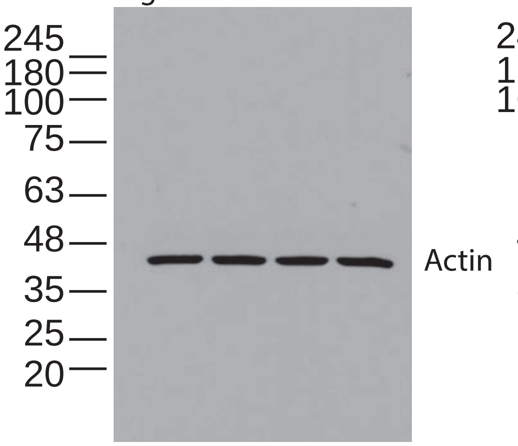


Fig 3B

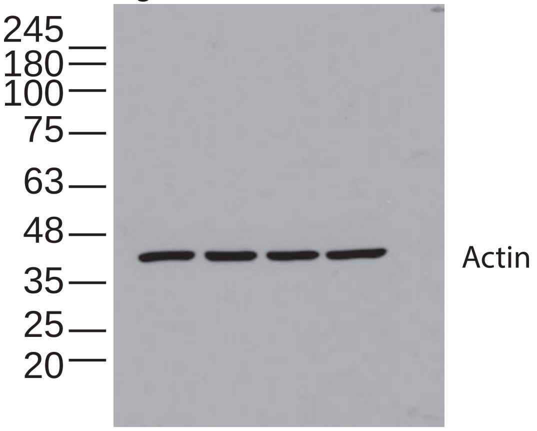

Fig 4A

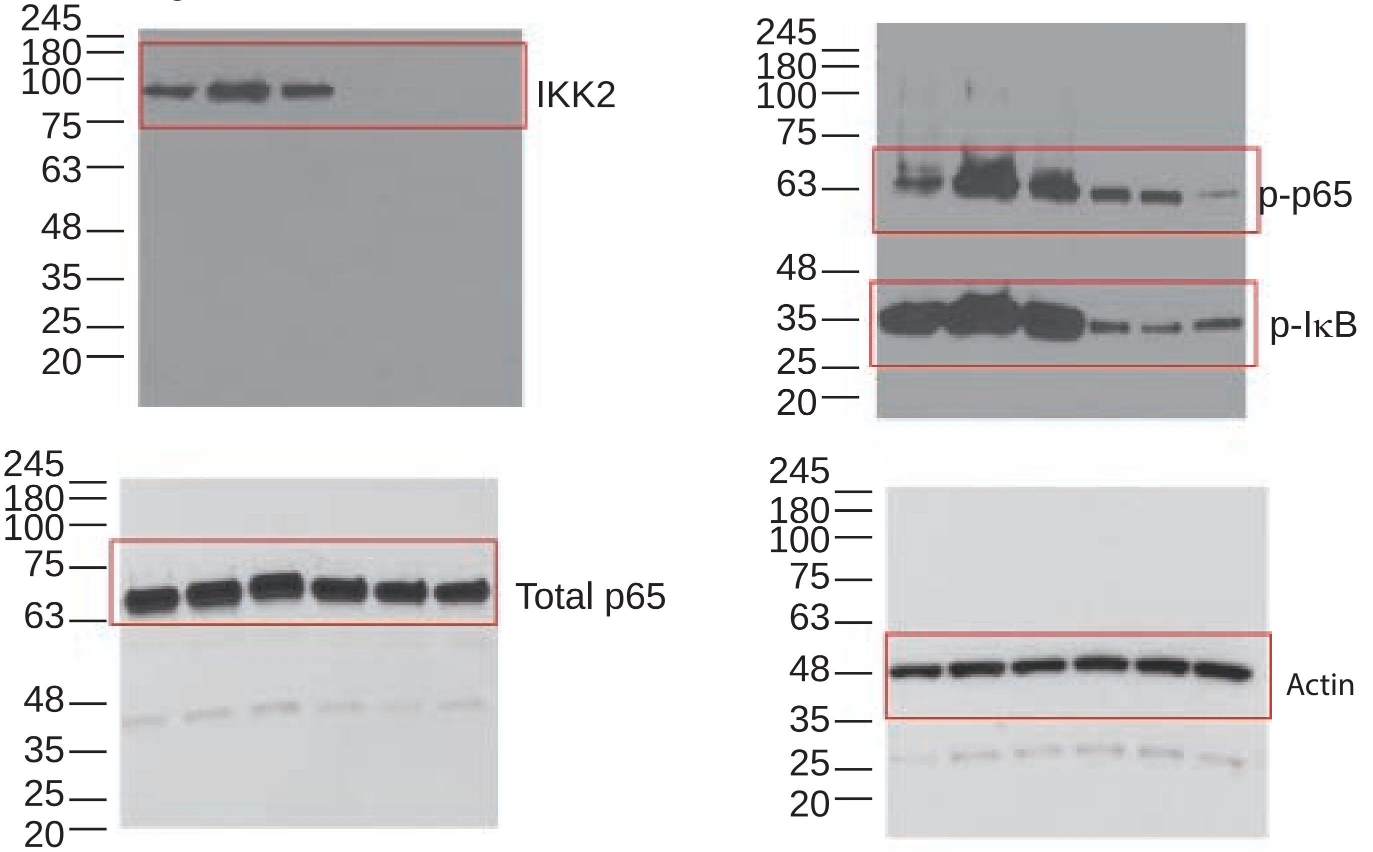


Fig 5A

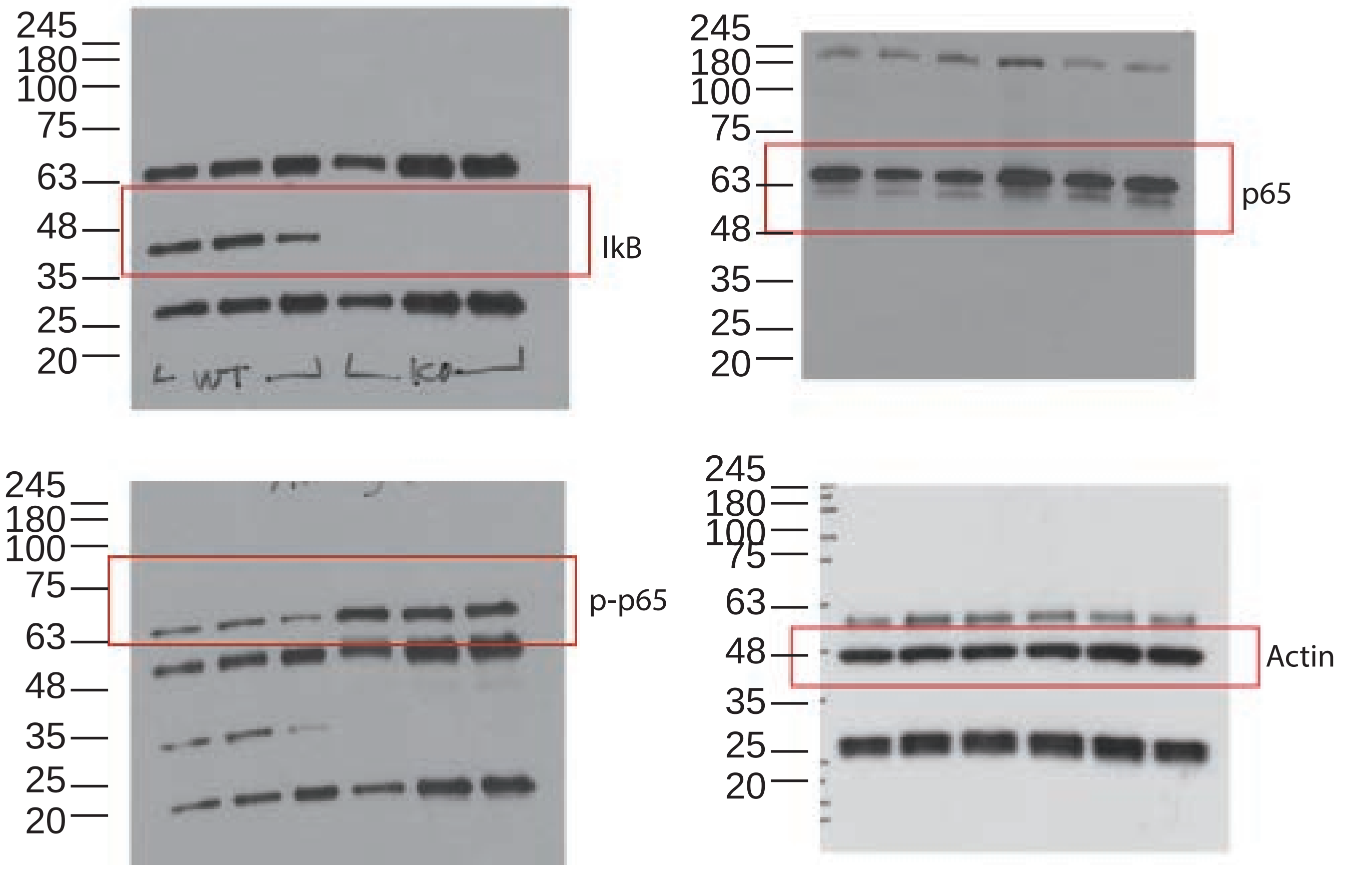

Fig 6A

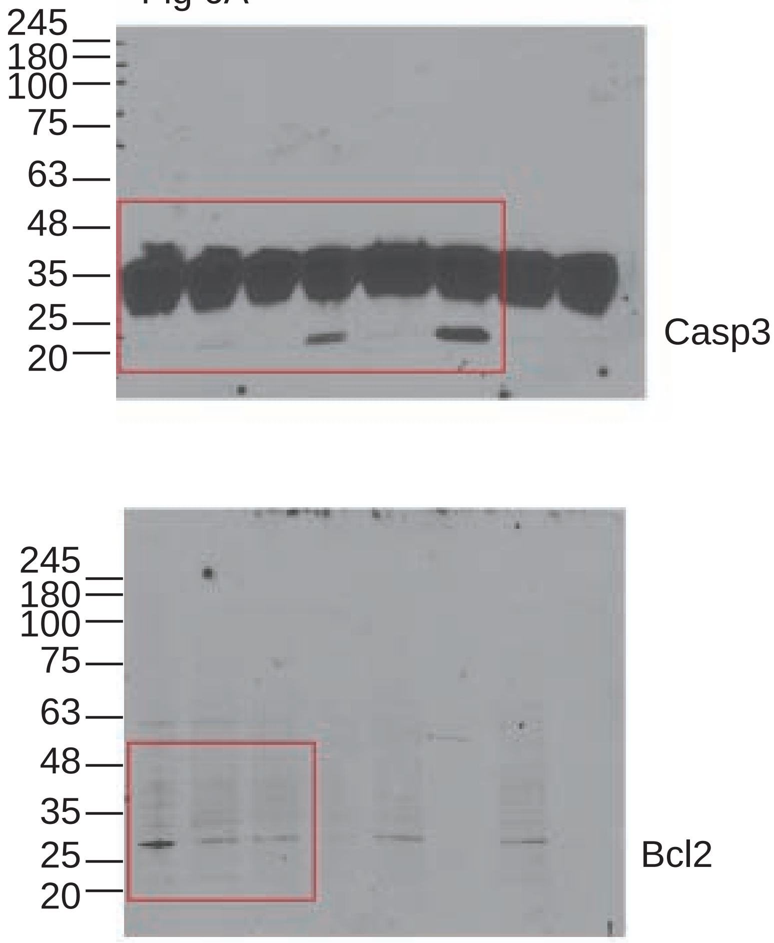


Fig 6B

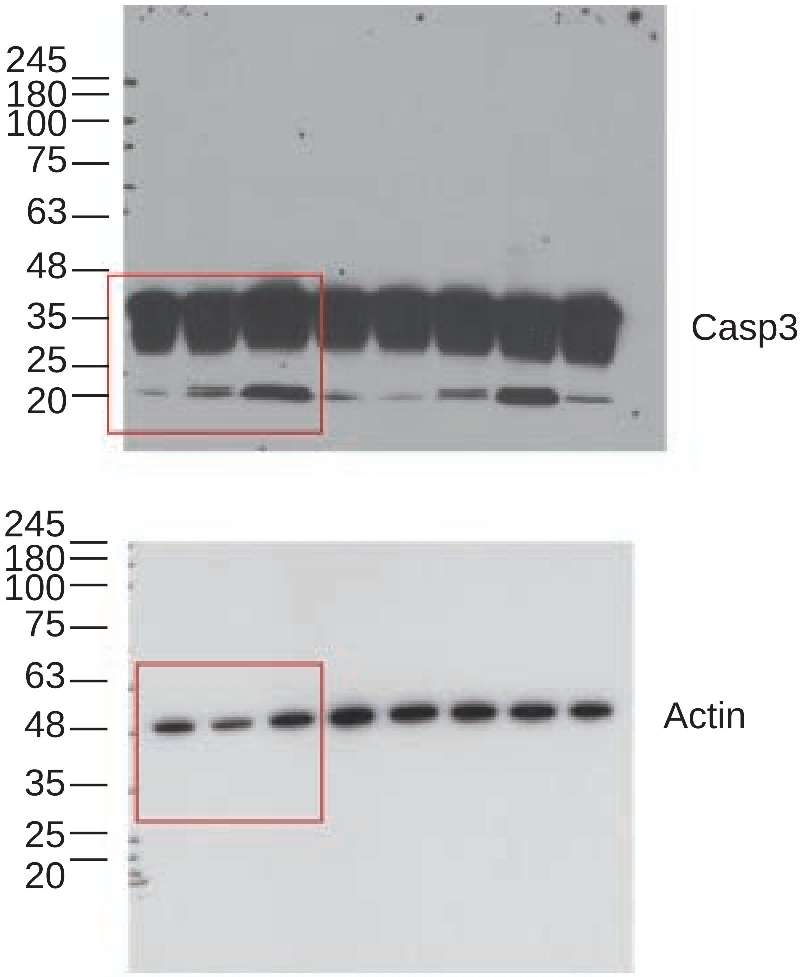


Fig 7F

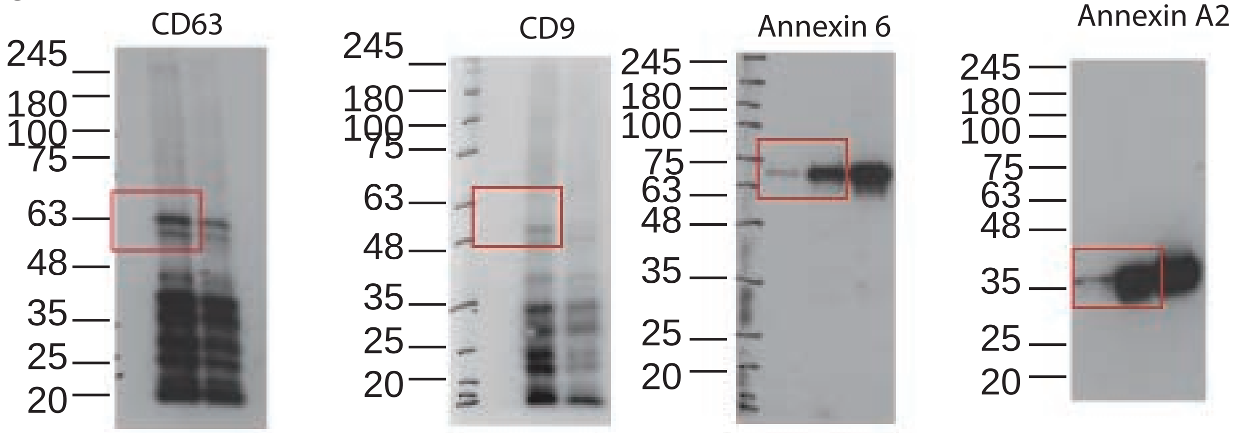


Fig 7J

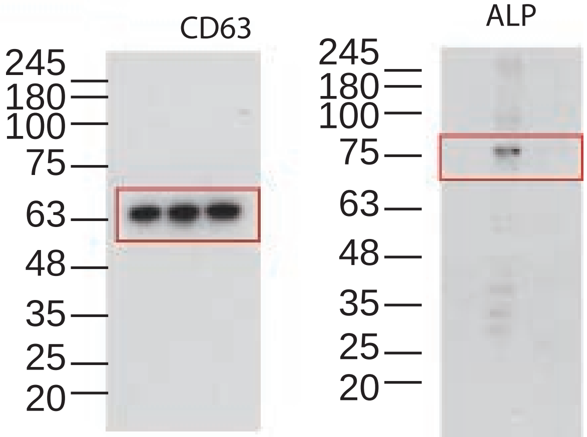
